# Deciphering cross-omics complexity of tissues via diagonal integration of unpaired spatial multi-omics data

**DOI:** 10.64898/2026.06.09.730286

**Authors:** Xiang Zhou, Kangning Dong, Jiankai Xiao, Luonan Chen, Shihua Zhang

## Abstract

Recent spatial multi-omics technologies enable the simultaneous *in situ* profiling of multiple omics modalities on the same tissue section; however, they face challenges in experimental complexity and high costs. This technical limitation can be circumvented by diagonal integration methods, which integrate omics data from different modalities. However, existing single-cell diagonal integration approaches overlook spatial information, causing unreliable anchoring across omics layers. Here, we introduce STAMO, a graph attention neural network model for spatially aware integration of unpaired spatial slices from different omics. Systematic benchmarking on spatial epigenome-transcriptome slices proves that STAMO outperforms the state-of-the-art methods in generating aligned embeddings and identifying consensus spatial domains across omics. We apply STAMO to integrate unpaired data from diverse spatial omics types (transcripts, epigenetics, DNA, and proteins), including slices from spatial RNA and four different epigenomic modalities, spatial ATAC and RNA slices across embryonic stages, spatial protein and RNA slices, and spatial DNA and RNA slices. In addition, the integration capability of STAMO can be further used to achieve cross-omics generation, offering a solution for exploring spatial region-specific gene regulatory mechanisms.

## Introduction

Rapid developments in spatial omics technologies have enabled the measurement of multiple omics layers, such as spatial chromatin accessibility (spATAC-seq^1^), histone modification (spCUT&Tag-seq^2^), DNA (slide-DNA-seq^3^), proteome (CODEX^4^), and transcriptome (10x Visium^5^), each offering unique insights into the state and function of complex tissue architecture. While spatial assays for the simultaneous profiling of multiple omics on the same section have recently been reported, such as spATAC-RNA-seq and spCUT&Tag-RNA-seq^6^, the co-profiling experiments still face technical trade-offs between sensitivity and throughput due to the complexity of the processing. A straightforward alternative approach is to measure different omics layers on separate or adjacent tissue sections, resulting in unpaired data, and then computationally integrate them to obtain in silico multi-omics data^3,7-9^.

In single-cell data analysis, integrating data from different omics faces diagonal integration issues^10^ where feature spaces across omics (e.g., genes, peaks, proteins) are mismatched. Integrating unpaired spatial slices brings additional challenges, including spatial location information modeling, variable spatial resolutions of different omics measurements (e.g., Visium and CODEX), and potential discrepancies between even closely adjacent slices. Existing methods for single-cell multi-omics integration^11-16^, such as Seurat v3^11^ and GLUE^12^, have made significant improvements in generating a unified joint representation for different omics data. However, these methods do not consider the modeling of spatial contexts, which hampers their performance on spatial omics data.

Existing spatially aware integration methods, such as CellCharter^17^, GraphST^18^, STAligner^19^, and PASTE^20^, focus on integrating spatial transcriptomics slices using shared genes. To apply these single-omics tools, multi-omics data need to be transformed into the same feature space based on prior knowledge. For example, spATAC-seq data can be converted into a gene activity matrix by counting DNA reads aligned near and within the gene body; however, such conversion results in information loss^21^. Integrating spatial RNA-seq and antibody-based proteomic data is also challenging, since the overlap between proteins and their corresponding protein-coding genes is limited, and a large number of omics-specific features are ignored by existing integration methods^18,19^. Recently proposed spatial multi-omics integration methods, SpatialGlue^22^ and SpaMosaic^23^, require known spot-paring information as a bridge to guide integration, lacking the ability to handle the general case of fully unpaired integration. Another spatial multi-omics integration strategy adopts image registration between adjacent tissue sections, such as spatial transcriptomics and proteomics^7,8^, spatial transcriptomics and metabolomics^9^. However, the inherent heterogeneity in tissue architecture between even closely adjacent sections makes it difficult to achieve precise spatial registration.

Cross-omics generation or translation, i.e., inferring one omics modality from another, also constitutes a critical aspect in multi-omics integration. While recent spatial-aware approaches such as SpaMosaic^23^ and NicheTrans^24^ have demonstrated strong performance in this task, they rely heavily on the availability of paired multi-omics data during model training. Consequently, these methods are inapplicable in scenarios where spatial multi-omics data are completely unpaired, highlighting an urgent need for integrative frameworks that can effectively perform cross-omics generation under unpaired conditions.

To this end, we introduce STAMO for spatially aware integration and generation for unpaired spatial multi-omics slices. STAMO adopts a variational graph attention autoencoder framework to effectively integrate spatial information, omics information, and biological prior knowledge. STAMO further introduces an adversarial learning strategy and optimal transport algorithm to correct the distribution differences of diverse omics. STAMO supports cross-omics data generation without paired information, saving the high cost of generating paired multi-omics data. Benchmarking of STAMO on multiple spatial epigenomic-transcriptome datasets demonstrated consistent outperformance in spot-spot matching, spatial domain identification, and cross-omics generation. STAMO can also decipher consistent tissue structures across omics slices, including spatial protein and RNA slices of human lymph node, and spatial DNA and RNA slices of mouse liver metastases.

## Results

### Overview of STAMO

The main idea of STAMO is to learn spot embedding containing spatial location information and feature embedding embedded with biological prior knowledge through graph neural networks, and then reconstruct the feature-by-spot count matrix data through dot multiplication (**Fig. 1a**). Here, spot is used as a shorthand to refer to either segmented cell or capture location (bead, bin, or spot), depending on the spatial omics platform employed. For each omics slice, spatial coordinates are used to build a spatial neighbor graph where nodes represent spots and edges indicate spatial proximity.

**Fig. 1.**
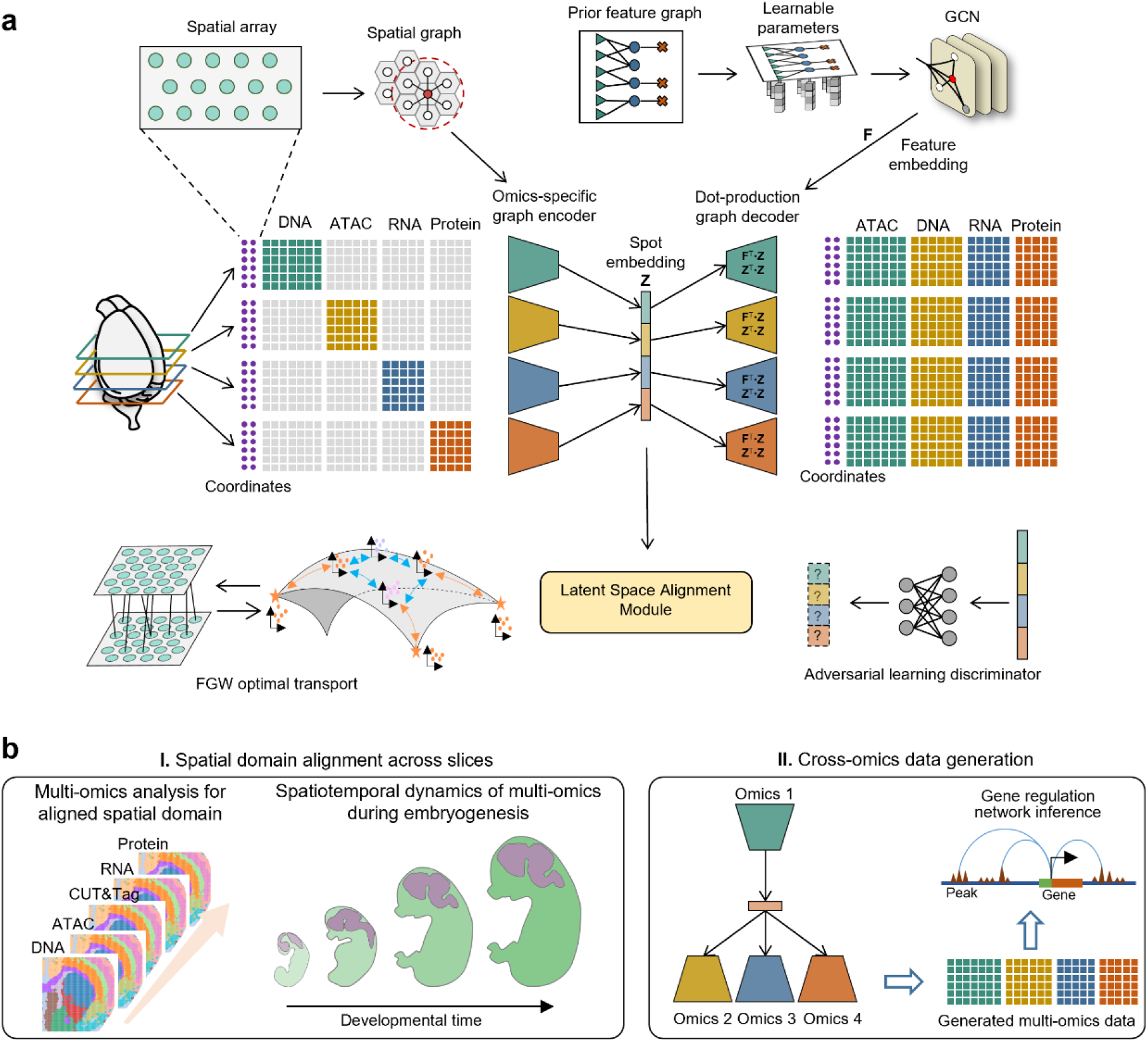
The overview of STAMO. **a**. STAMO adopts a two-stage training strategy. In stage 1, pretrain a graph attention network to produce coarse-aligned embeddings. In stage 2, identify anchors via Fused Gromov-Wasserstein optimal transport and perform anchor-guided alignment. **b**. STAMO outputs integrated spot representations and the trained model enables multi-omics data generation from profiled single-omics slices. The results can be used to identify consensus spatial domains across unpaired omics slices with distinct omics feature spaces (including DNA, CUT&Tag, ATAC, RNA, and Protein), slices from different developmental stages (**I**) and gene regulation network inference (**II**).

For the spot embedding processing, STAMO adopts an omics-specific variational graph attention encoder to learn spatially aware embeddings for each spot by aggregating information from neighboring nodes. Unlike existing ST integration methods via shared autoencoder^18,19^, STAMO employs a separate autoencoder for each omics layer to model the distinct distributions of different omics data. For example, coupled with the prior information of proteins and their associated coding genes, STAMO can independently model protein abundances and gene expressions. Additionally, to mitigate cross-omics and cross-slice heterogeneity, both adversarial learning strategy and Fused Gromov-Wasserstein (FGW) optimal transport (OT) algorithm are introduced here.

For feature embedding processing, we first construct a prior feature graph, and then leverage the graph convolutional neural network with trainable node features to obtain its low-dimensional representation. Specifically, the feature graph is constructed based on chromatin distance between genes and peaks, or protein-gene coding relationship.

Once trained, STAMO produces integrated latent embeddings that support joint analysis across different omics slices with distinct feature space (e.g., DNA, CUT&Tag, ATAC, RNA, and protein) and slices from different developmental stages, enabling consensus tissue structures identification (**Fig. 1b**). Beyond alignment, the model further supports cross-omics data generation by encoding data from one omics into the integrated latent space and decoding it into another. This generative capability allows STAMO to computationally reconstruct missing omics, thereby facilitating the investigation of spatial region-specific gene regulatory mechanisms.

### Benchmarking on diagonal integration and cross-omics data generation performance on the spatial transcriptome-epigenome dataset

To quantitatively evaluate the diagonal integration performance of STAMO, we applied it to four spatial transcriptome-epigenome coronal brain datasets of postnatal day 22 (P22) mice with known spot-to-spot pairing information^6^. The data include one spATAC-RNA-seq slice and three spCUT&Tag-RNA-seq slices (H3K27ac, H3K27me3, and H3K4me3). To simulate a scenario for unpaired multi-omics slices integration (**Fig. 2a**), we removed the spot-pairing information and constructed four RNA-epigenome modality pairs: RNA + H3K27ac, RNA + H3K27me3, RNA + H3K4me3, and RNA + ATAC. For benchmarking, we compared STAMO against both non-spatial single-cell multi-omics integration methods—Seruat, scVI, and GLUE—as well as the spatially aware spatial transcriptomics integration method CellCharter and STAligner.

**Fig. 2.**
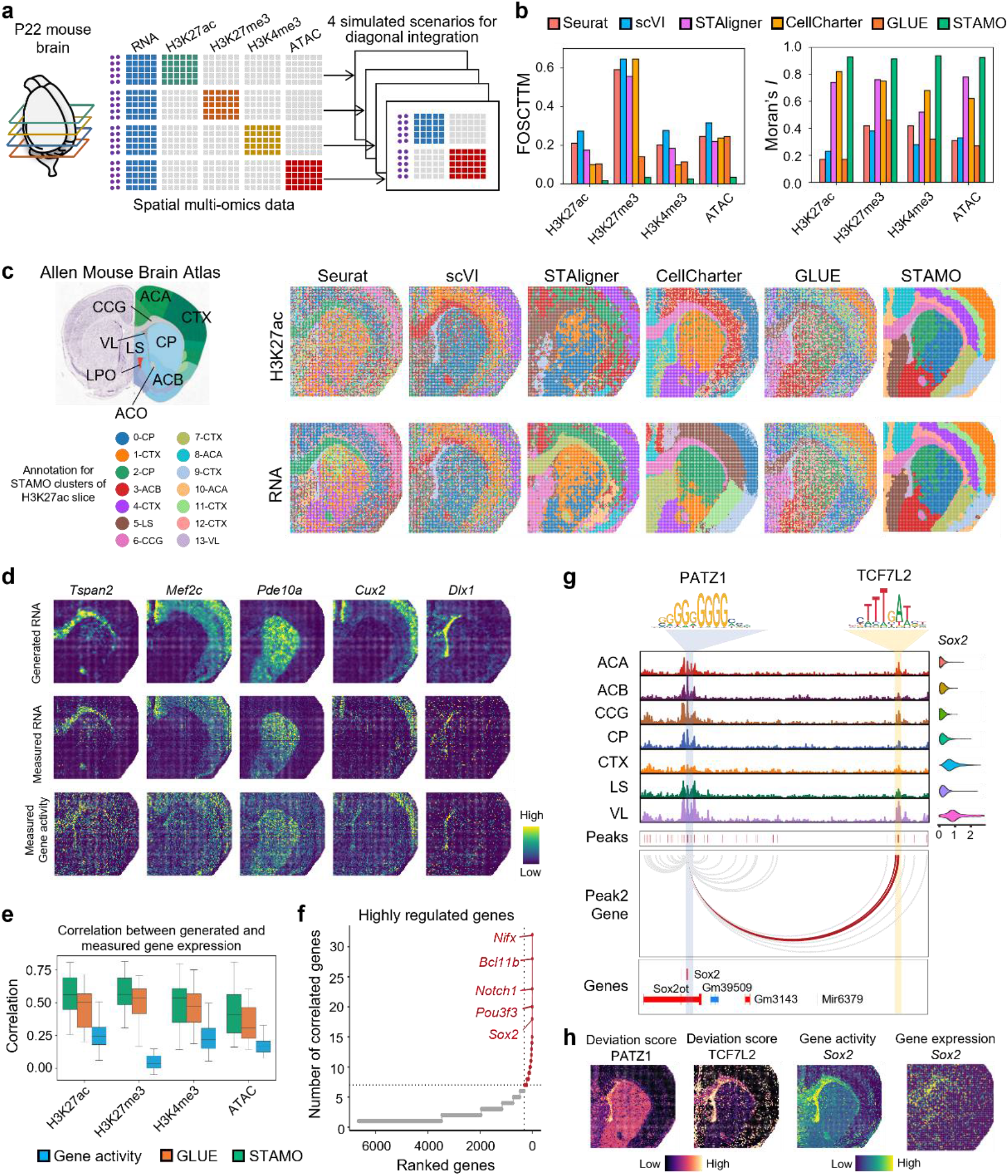
Benchmarking on diagonal integration and cross-omics data generation performance on the spatial transcriptome-epigenome dataset. **a**. Four P22 mouse brain spatial spCUT&Tag-RNA-seq and ATAC-RNA slices. To simulate a diagonal integration scenario, pairing information is masked. **b**. Spot level alignment error (quantified by FOSCTTM) (left) and Moran’s *I* score (right) of different integration methods. **c**. Allen Brain Atlas reference annotation for the postnatal mouse brain (left), Spatial domains characterized by different integration methods (right). The annotated clusters correspond to STAMO’s results, and the domain colors do not necessarily match the results of the other methods. **d**. Spatial heatmap of generated and measured gene expression for marker genes. **e**. Gene-wise correlation between the generated and measured gene expression, comparing STAMO with competing methods. **f**. Highly regulated genes based on the number of significant peak-gene links. Genes with >7 links are marked by a dashed line. **g**. Genome track visualization of the marker gene *Sox2* with peak-gene links between regulatory elements and the target gene. **h**. Spatial mapping of deviation score^104^ for TF PATZ1 and TCF7L2, gene activity, and gene expression for the target gene *Sox2*.

We first evaluated single-spot-level alignment accuracy via the FOSCTTM metric^25^, which quantifies how closely paired spots from different omics modalities are positioned in the integrated latent space. As shown in **Fig. 2b**, STAMO achieved the lowest FOSCTTM scores across all four datasets, highlighting its superior modality alignment performance. These results were further supported by the uniform manifold approximation and projection (UMAP) visualizations of the aligned embeddings, where STAMO exhibited well-mixed spots across slices (**Supplementary Fig. 1a-d**). In terms of spatial coherence, measured by Moran’s *I*—an indicator of spatial autocorrelation within identified clusters—STAMO consistently outperformed all other methods across the datasets. While GLUE ranked second in FOSCTTM, its Moran’s *I* scores were lower than those of the spatially aware method STAligner and CellCharter, highlighting the importance of incorporating spatial information in integration. Moreover, STAMO consistently outperformed competing methods in both biological conservation and batch mixing metrics, underscoring its strong capability for effective batch correction while preserving biologically meaningful embeddings (**Supplementary Fig. 2**). The spatial plots in the spatial domain further confirm these quantitative results. For example, in the integration of the H3K27ac and RNA slices (**Fig. 2c, right**), we found that STAMO’s spatial clusters had the least noise and greater cross-omics structural consistency compared to competing methods. Using the Allen Brain Atlas for annotation (**Fig. 2c, left**), we identified major brain structures corresponding to STAMO-derived clusters: cortex layers (CTX; cluster 1, 6, 7, 8, 11 and 12), genu of corpus callosum (CCG; clusters 4), lateral ventricle (VL; clusters 13), caudoputamen (CP; cluster 0 and 3), nucleus accumbens (ACB; cluster 2), lateral septal complex (LS; cluster 5). In contrast, non-spatial methods such as GLUE, scVI, and Seurat produced clusters with poorly defined boundaries. Although STAligner successfully captured CTX and CCG, it failed to consistently align CCG across slices. Similar spatial clustering improvements by STAMO were observed in the remaining three datasets (**Supplementary Fig. 3a–c**). To further validate the cluster annotation results, we visualized the gene expression of known marker genes for five major brain regions across all datasets (**Supplementary Fig. 4a–d**). Marker genes—including *Pde10a* for CP^26^, *Mef2c*^27^ and *Cux2*^28^ for CTX, *Tspan2*^29^ for CCG, and *Dlx1*^30^ for VL—showed distinct spatial patterns that were consistent with STAMO-derived clusters.

To clarify the contribution of each key component in STAMO architecture, we have performed three ablation experiments to evaluate the effects of the graph attention-based autoencoder (GAT), FGW-OT, and adversarial learning on integration performance. As shown in **Supplementary Figs. 5-6**, disabling any single component leads to a decline in integration performance. Notably, spatial information exerts the greatest impact on overall performance. Removing the GAT module severely compromises spatial domain detection and spot matching accuracy, with clear deteriorations in FOSCTTM, Moran’s *I*, and biological conservation scores. These results underscore the necessity of incorporating spatial context. Furthermore, ablating either the FGW-OT or the adversarial learning module substantially reduces batch mixing scores, indicating that both components are essential for generating modality-aligned embeddings. In the first training stage, adversarial learning produces an initial globally aligned embedding, which facilitates anchor point identification for the subsequent stage. In the second stage, the FGW-OT alignment module further refines the co-embedding to achieve local alignment.

Beyond cross-omics consensus spatial domain identification, the trained model was further used to generate missing data modalities. We evaluated its performance in predicting gene expression profiles from epigenomic slices. Among the compared methods, only GLUE is capable of cross-omics generation in the absence of paired multi-omics data, and was thus included as a baseline alongside gene activity scores derived from ATAC or CUT&Tag data. The generated profiles reconstructed by STAMO closely recapitulated the measured gene expression in spatial patterns of marker genes and exhibited stronger signals compared to gene activity scores (**Fig. 2d**). The results of the Pearson correlation coefficient between the generated and measured gene expression further confirmed that STAMO achieved higher generation accuracy than competing methods (**Fig. 2e**). We believe this is due to its explicit modeling of spatial information in the generation, which enables neighborhood-level signal fusion and enhancement. This result also highlights the limitation of integration methods that rely on gene activity scores (such as Seruat, scVI, CellCharter and STAligner), which did not perform well, especially in H3K27me3 data with transcriptional repression.

We also assessed whether the generated data preserves gene-gene correlation structure (e.g., co-expression networks). Specifically, we computed the gene-gene correlation matrices for both generated and measured gene expression data, and quantified their similarity using Spearman correlation. As shown in **Supplementary Fig. 7a**, STAMO achieves the highest average correlation of 0.83 among all competing methods, indicating superior preservation of gene-gene relationships. We next examined the utility of generated data in downstream analyses, including spatial domain identification (clustering) and differential expression analysis. As shown in **Supplementary Fig. 7b**, STAMO demonstrates the strongest clustering consistency compared with other methods. Moreover, the log fold-change (logFC) values derived from generated and measured data show strong agreement, with average Spearman correlations of 0.97 for high-confidence genes (*q*-value < 0.01) and 0.79 for all genes (**Supplementary Fig. 7c**), further supporting the biological fidelity of the generated data.

To further validate the biological relevance of the generated data, we examined the correlation between the generated gene expression and gene activity scores (GAS) or chromatin silencing scores (CSS)^6^ within specific anatomical regions. For example, in the CCG region (**Supplementary Fig. 8a**), we observed a robust anticorrelation between the generated gene expression and measured H3K27me3 CSS, and positive correlations between the generated gene expression and measured GAS derived from H3K27ac, H3K4me3, and ATAC data. We found that genes such as *Grin2b* with high CSS and low RNA expression were related to synaptic transmission and neurotransmitter release^31^. Genes such as *Mal* and *Mag* with high GAS and high RNA expression were related to myelination and regulation of oligodendrocyte differentiation, which were in accordance with previous findings^6^. These results give confidence in the accuracy of the generated data.

Using data generated by STAMO, we could computationally construct paired multi-omics profiles from single-omics slices, thereby enabling the identification of peak-gene regulatory links. As a demonstration, we focused on the H3K27ac assay for peak-gene correlation analysis using the FigR framework^32^, prioritizing genes based on the number of significantly correlated peaks (**Fig. 2f**). This analysis successfully recovered known marker genes, including *Sox2*^*33*^, *Notch1*^*34,35*^, and *Nfix*^*36*^ for the VL region, as well as *Bcl11b* for the CP region^6^. We further investigated candidate transcription factor (TF) regulators for *Sox2*, a key gene involved in neural development and stem cell maintenance^37,38^ (**Supplementary Fig. 8b**). Notably, binding motifs for *Patz1*^*39,40*^ and *Tcf7l2*^*41,42*^ were located within the promoter and enhancer regions of *Sox2*, respectively, suggesting distinct regulatory roles in modulating *Sox2* expression within the VL region^6^ (**Fig. 2g**). The spatial co-localization of TF motif enrichment (*Patz1* and *Tcf7l2*), gene activity (*Sox2*), and *Sox2* gene expression further supported these regulatory relationships (**Fig. 2h**). Collectively, these results demonstrate STAMO’s capability to uncover key regulatory elements through the integration of unpaired spatial multi-omics datasets.

### Five-omics diagonal integration enables simultaneously analysis of both positive and negative epigenetic regulation

ATAC, H3K27ac, and H3K4me3 generally mark active chromatin regions and are associated with transcriptional activation, whereas H3K27me3 is typically linked to gene repression, marking regions where gene expression is silenced. To jointly investigate the interplay between these epigenetic modalities and gene expression, we applied STAMO to integrate the four P22 mouse brain datasets^6^ encompassing five distinct omics layers: transcriptome, chromatin accessibility, and three histone modifications. Currently, no spatial multi-omics technology can simultaneously profile all five omics layers. To simulate a realistic diagonal integration scenario, we selected ATAC, H3K27ac, H3K27me3, and H3K4me3 slices without corresponding RNA measurements and RNA slices from the spH3K27ac-RNA-seq slice, removing spot-pairing information. Thus, we can obtain five unpaired slices from different omics modalities (**Fig. 3a**).

**Fig. 3.**
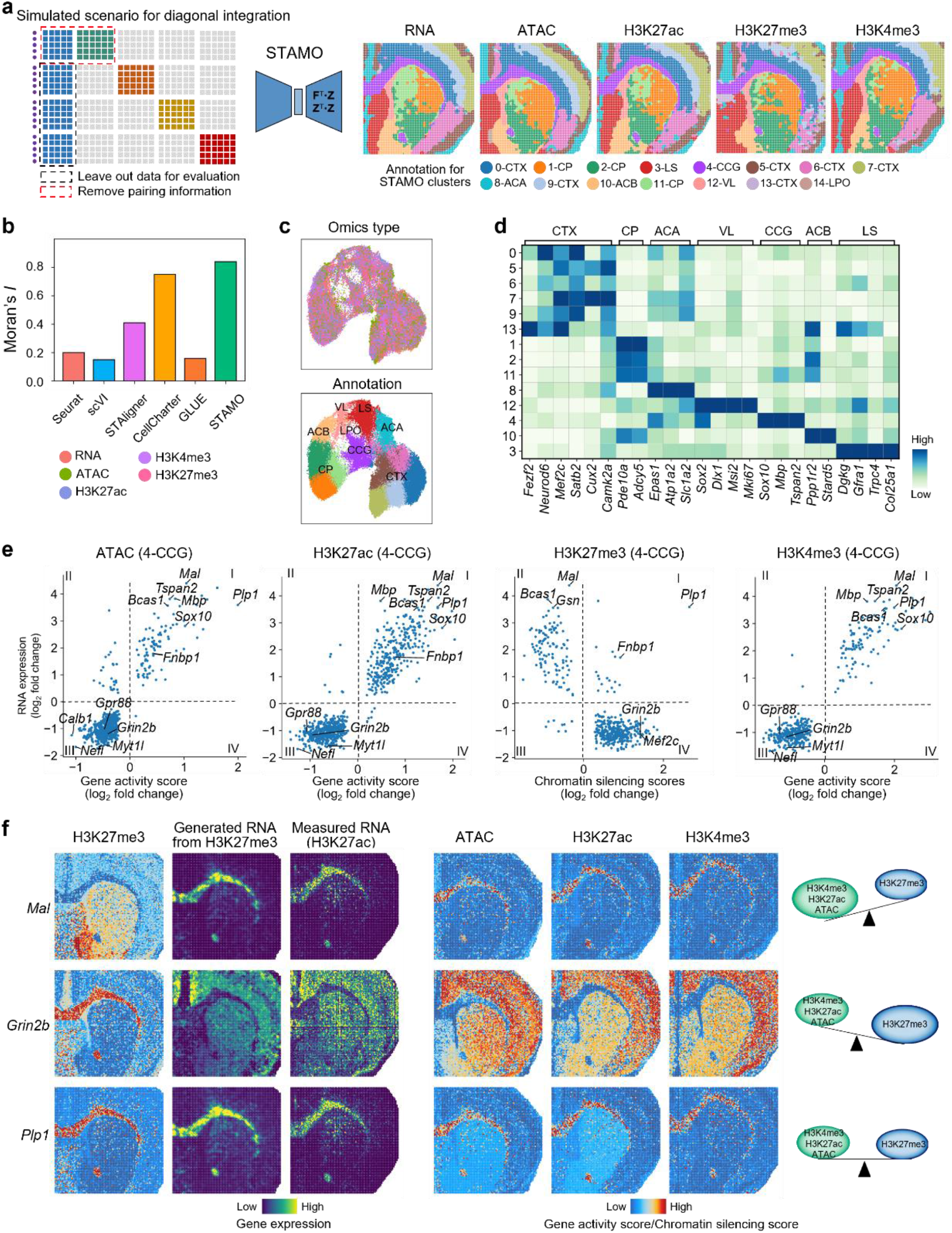
Five-omics diagonal integration enables simultaneously analysis of both positive and negative epigenetic regulation. **a**. Four P22 mouse brain spatial spCUT&Tag-RNA-seq and ATAC-RNA slices. To simulate a five-omics diagonal integration scenario, all pairing information was removed. RNA slices from spH3K27ac-RNA-seq data were used for training, while other RNA slices were excluded for evaluating cross-omics data generation (left). Spatial domains characterized by STAMO (right). **b**. Moran’s *I* score of different integration methods. **c**. UMAP plots of the embeddings from STAMO. Spots are colored by omics type (top) and colored by cluster (bottom). **d**. Heatmap of marker gene expression for the identified spatial domains by STAMO. **e**. Correlation between ATAC GAS, H3K27ac GAS, H3K27me3 CSS, H3K4me3 GAS, and RNA gene expression of the identified CCG region. **f**. Spatial heatmap of generated and measured RNA gene expression, GAS/CSS for CCG-specific marker genes *Mal, Grin2b*, and *Plp1* with different gene regulation patterns.

We first evaluated the joint clustering results produced by different integration methods (**Fig. 3a, Supplementary Fig. 9a**). Consistent with previous observations, STAMO generated cleaner spatial clusters than competing methods, which was quantitatively confirmed by higher Moran’s *I* score and biological conservation scores (**Fig. 3b, Supplementary Fig. 9b**). The identified clusters corresponded well to known brain regions as annotated in the Allen Brain Atlas. For instance, STAMO accurately delineated the CTX (clusters 0, 5, 6, 7, 9, and 13), CCG (cluster 4), CP (clusters 1, 2, and 11), ACB (cluster 10), and VL (cluster 12). In addition to capturing cross-omics shared spatial domains, STAMO also identified slice-specific structures such as the lateral preoptic area (LPO; cluster 14), uniquely present in the ATAC slice, highlighting its ability to detect slice-specific spatial structures. We also used UMAP visualization to visualize the data before and after integration. Prior to integration, larger modality gaps were observed between RNA and H3K27me3 compared to the other omics (**Supplementary Fig. 9c**),underscoring the need for integration methods that are flexible and robust to cross-omics heterogeneity. After STAMO integration, the data from different omics slices were well mixed while preserving clear spatial domain separation (**Fig. 3c**). The annotations were further validated by elevated expression of known marker genes, such as *Mbp* and *Tspan2* for the CCG, and *Fezf2* and *Cux2* for deep and superficial cortical layers, respectively^6^ (**Fig. 3d**).

With the unified spatial domains established across the five omics layers, we next explored region-specific regulatory mechanisms underlying gene expression. Taking the CCG region as an example, we observed strong positive correlations between ATAC, H3K27ac, H3K4me3 and gene expression, but H3K27me3 was inversely correlated (**Fig. 3e**). *Mal*, a marker gene in CCG, showed enrichment in ATAC, H3K27ac, and H3K4me3, with minimal enrichment in H3K27me3 (**Fig. 3f**). Conversely, *Grin2b* was repressed in CCG, with high H3K27me3 enrichment, illustrating the mutually exclusive regulatory relationship between activating (ATAC/H3K27ac/H3K4me3) and repressive (H3K27me3) epigenetic states. In addition, we identified instances of co-occupancy of activating and repressive marks (known as bivalent chromatin states^43^) at several loci, such as *Plp1*^44^, which may reflect transcriptional poising. We also benchmarked the generation performance of STAMO in reconstructing gene expression profiles for the four epigenomic slices. As shown in **Fig. 3f** and **Supplementary Fig. S9d-e**, STAMO again achieved the highest generation performance and accurately generated gene expression under different epigenetic regulation roles (positive, negative, and co-occupancy). Collectively, these results demonstrate STAMO’s capacity to integrate diverse omics datasets and elucidate complex, region-specific epigenetic regulatory mechanisms.

### Integrating mouse embryo brain slices across different developmental stages reveals spatiotemporal dynamics of embryogenesis

Beyond cross-omics diagonal integration, STAMO is also capable of performing cross-batch integration by correcting batch effects. To evaluate its performance, we applied STAMO to a mouse embryonic brain dataset generated by MISAR-seq^45^, which jointly profiles gene expression and chromatin accessibility (spATAC-RNA). The dataset spans four developmental stages (E11.0, E13.5, E15.5, and E18.5), each consisting of a paired spATAC-RNA slice. To simulate diagonal integration scenarios, we removed the spot-pairing information from each slice following the previous procedure (**Fig. 4a**). As shown by UMAP visualization, substantial omics differences and strong batch effects were observed across developmental stages (**Supplementary Fig. 10a**). The manual annotations and spot-pairing information were used as ground truth for benchmarking.

**Fig. 4.**
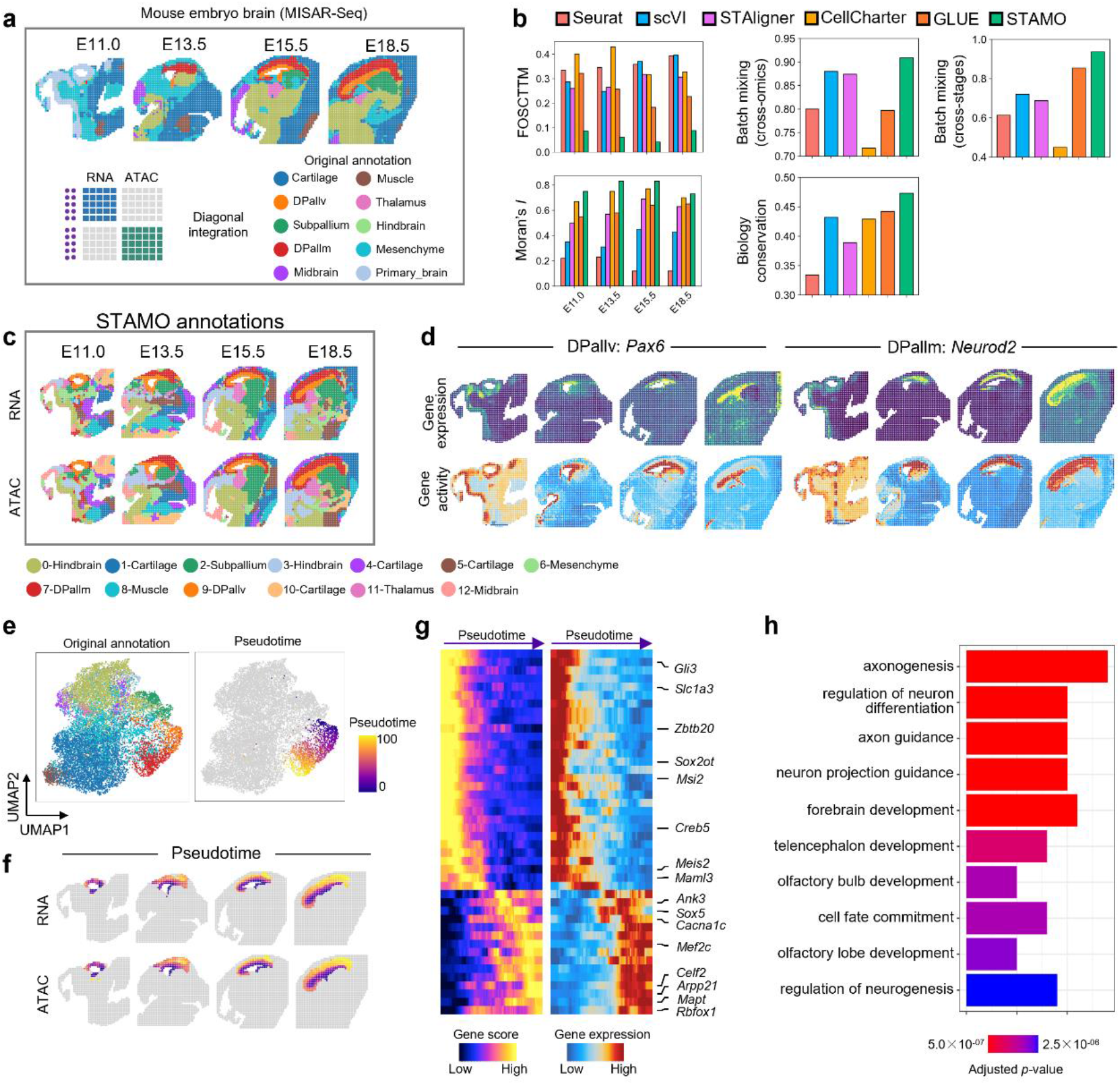
Integrating mouse embryo brain slices across different developmental stages reveals spatiotemporal dynamics of embryogenesis. **a**. Four spatial ATAC-RNA slices produced by MISAR-Seq (E11.0 to E18.5). Pairing information was masked during integration. **b**. Spot-level cross-omics alignment error (FOSCTTM, left) and biological conservation versus batch mixing scores (right) of different integration methods. **c**. Spatial domains characterized by STAMO. **d**. Spatial visualization of marker genes relating to the domain Dpallm and Dpallv identified by STAMO. **e**. UMAP plots of embeddings of STAMO colored by original tissue annotations (left) and the trajectory of corticogenesis (right). **f**. Spatial mapping of the pseudotime of corticogenesis. **g**. Heatmap showing gene activity score and gene expression along pseudotime. **h**. GO enrichment on the top variable genes along the pseudotime. The top ten significant GO terms are shown.

STAMO achieved the best cross-omics spot alignment performance with the lowest FOSCTTM scores in all four time points (**Fig. 4b**). Moreover, STAMO outperformed other methods in both biological conservation and batch mixing metrics, highlighting its effectiveness for simultaneous cross-omics and cross-batch integration. These results were consistent with the UMAP visualizations (**Supplementary Fig. 10a**), where data from different omics and stages were well mixed, while spatial domains remained clearly separated. Compared to competing methods, STAMO produced spatial clusters with least noise and highest concordance with the original annotations (**Fig. 4c**). It not only accurately resolved large tissue structures such as cartilage (clusters 1, 4, 5, 10), mesenchyme (cluster 6), and hindbrain (clusters 0, 3), but also successfully identified rare structures such as muscle (cluster 8), thalamus (cluster 11), midbrain (cluster 12), dorsal pallium ventricular zone (DPallv, cluster 9), and dorsal pallium mantle zone (DPallm, cluster 7). In contrast, competing methods yielded noisier outputs and could only roughly differentiate several major categories with significant biological differences (**Fig. 4c, Supplementary Fig. 10b**).

We further examined clusters 7 and 9 in the E13.5 slice, which overlapped with DPallm and DPallv regions. In the original annotation (**Fig. 4a**), DPallm localized to the upper and right regions surrounding the central cavity, while DPallv was located below it. In STAMO’s results, cluster 7 corresponded well to DPallm, while cluster 9 encircled the cavity, exhibiting a slightly different spatial pattern compared to the original annotation. We validated the assignment of cluster 9 by the expression of *Pax6*, a key regulator of mouse neocortex development, which is highly expressed in DPallv^46^. High gene expression and gene activity scores in cluster 9 confirmed the annotation (**Fig. 4d**).

Importantly, STAMO identified spatial domains that remained highly consistent not only across omics layers but also across developmental stages. We observed increasing proportions of DPallm, hindbrain, and subpallium structures (**Supplementary Fig. 11**), reflecting the proliferation of these tissue structures during organogenesis^47,48^. Notably, at the earliest stage, E11.0, clusters annotated as DPallm, DPallv, hindbrain, and midbrain show a high degree of overlap with the primary brain in manual anatomical annotations reported in the original study, suggesting potential developmental relationships between the primary brain and these structures^49,50^. This spatiotemporal trajectories across the four stages were further validated by the expression of region-specific marker genes^45^: *Snhg11* (hindbrain), *Neurod2* (DPallm), *Pax6* (DPallv), and *Rnf220* (thalamus) (**Fig. 4d, Supplementary Fig. 10c**). In contrast, all baseline methods struggled to maintain consistent spatial domains across omics layers and developmental stages. For example, in GLUE’s results, the DPallm region was assigned to different clusters between E13.5 and E15.5. STAligner was able to align DPallm across developmental stages but failed to maintain cross-omics consistency. Altogether, these results highlight STAMO’s superior performance in identifying complex tissue structures and accurately aligning them across both omics modalities and developmental stages.

Based on the integrated embedding, we can further infer consensus pseudotime across omics and developmental stages to investigate the spatiotemporal dynamics of accessibility activity and gene expression. We selected the cortex regions to analyze the corticogenesis trajectory. The pseudotime axis was found to have captured the temporal order (E11.0 to E18.5) and spatial inside-out trajectory from the ventricular zone (DPaIIv) to cortical plate (DPallm) (**Fig. 4e-f**). We further conducted GO analysis on the top variable genes along the pseudotime, and found significant enrichment in functions including axonogenesis, regulation of neuron differentiation, forebrain development, cell fate commitment, and regulation of neurogenesis^51-66^ (**Fig. 4g-h**). These results suggest that these genes are likely involved in driving cell fate decision and establishing the inside-out patterning of cortex.

### Integration of spatial transcriptomics and proteomics slices with limited overlap features

Integrating spatial transcriptomics and proteomics offers a more comprehensive understanding of tissue complexity. We applied STAMO to integrate two human lymph node slices profiled by the 10x Xenium spatial transcriptomics platform with a 377-gene panel, and the CODEX spatial proteomics platform with a 28-marker antibody pane^l67^ (**Fig. 5a**). To link features across datasets, protein names were manually converted to their corresponding coding gene names based on established protein-gene relationships. After conversion, only 15 shared features remained, posing a significant challenge for existing integration methods that adopt shared features as input. Despite this, STAMO successfully integrated the two datasets. UMAP visualization of the aligned embeddings demonstrated well-mixed spots across slices while preserving clear spatial domain separations characterized by distinct expression patterns (**Supplementary Fig. 12a-b**). We manually annotated each spatial domain according to the known functions of differentially expressed genes. STAMO correctly identified canonical lymph node structures across RNA and protein slices (**Fig. 5b, Supplementary Fig. 12c**)^67^, including follicle region enriched with B cells highly expressing gene *MS4A1* and its encoded protein CD20^68^, paracortex region enriched with T cells highly expressing gene *CD3E* and its protein product CD3E^69^, medulla region highly expressing macrophage marker genes *CD163*^*70*^ and *CD68*^*71*^, along with lymphatic vessels marker *LYVE1*^*72*^ and their corresponding proteins CD163, CD68, and LYVE1, respectively. Small vascular structures were also identified, confirmed by the elevated expression of the smooth muscle marker *ACTA2*^*73*^ and its protein SMACTIN.

**Fig. 5.**
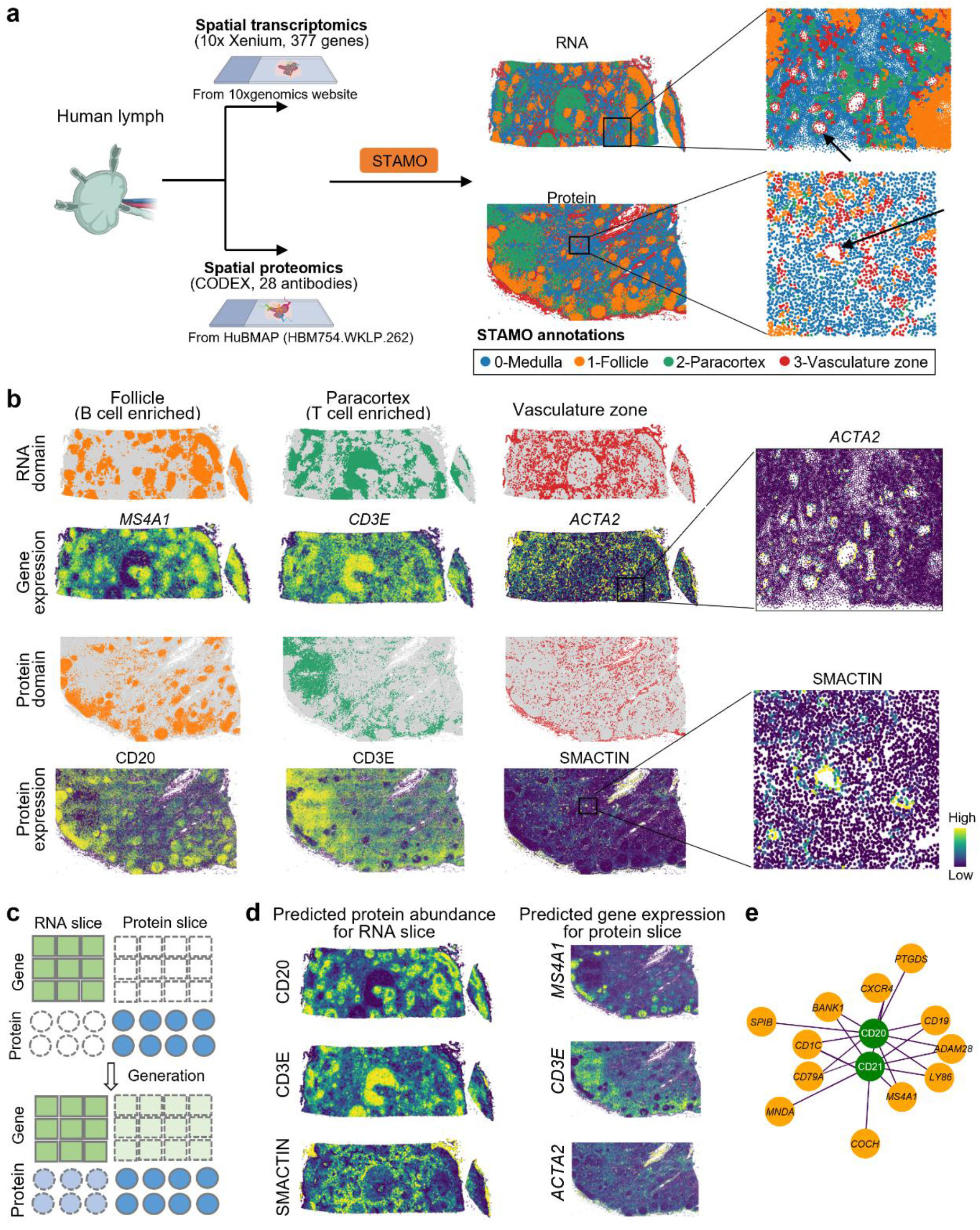
Integration of human lymph spatial transcriptomics and proteomics slices with limited overlap features. **a**. Human lymph spatial transcriptomics slice produced by 10x Xenium and spatial proteomics slice produced by CODEX (left). Visualization of identified spatial domains across slices characterized by STAMO (right). Created in https://BioRender.com. **b**. Spatial visualization of annotated tissue structures showing marker gene expression (top) and corresponding protein expression (bottom). **c**. The cross-omics generation schematic for protein and RNA slices. **d**. Spatial heatmap of marker protein abundance (left) and gene expression (right) predicted by STAMO. **e**. Gene-protein correlation network inferred using STAMO’s predicted protein data and measured RNA data, centered at B cell marker protein CD20 and CD21.

STAMO further enabled cross-omics prediction of missing data modalities, generating gene expression profiles for the CODEX protein slice and protein abundances for the Xenium RNA slice (**Fig. 5c**). The predicted protein abundance closely matched the measured gene expression spatial patterns in the RNA slice (**Fig. 5d**). For example, the predicted CD20 abundance accurately captured spatial specificity of lymphatic follicle with spherical or ovoid structures, and was localized in *MS4A1*-enriched B cell regions. Similarly, predicted gene expression on the CODEX slice reflected consistent spatial patterns with corresponding protein measurements. Leveraging the predicted protein abundance in the Xenium RNA slice, we inferred gene-protein correlation networks. Using B cell-enriched proteins CD20^68^ and CD21^74^ as examples, we visualized the top 10 most correlated genes (**Fig. 5e**). CD20 was strongly correlated not only with its coding gene *MS4A1*, but also with multiple canonical B cell markers such as *BANK1*^*75*^, *CD19*^*76*^, *CD79A*^*77*^, and *CXCR4*^*78*^, which also showed correlations with CD21.

Integrating data from Xenium and CODEX slices is relatively straightforward, as both platforms are subcellular resolution. We further evaluated STAMO’s performance in a more challenging scenario by integrating the subcellular resolution CODEX slice with a 10x Visium human lymph node slice at spot-level resolution, where each spot represents multiple cells (**Supplementary Fig. 13a-b**). The CODEX platform features immunofluorescent imaging of immune, lymphatic, and blood vessel markers^67^, enabling the identification of major cell types. We can find that main lymph node architectures were identified across slices. UMAP visualization again confirmed successful integration, revealing well-mixed embeddings and clear spatial domains separation defined by RNA and protein markers (**Supplementary Fig. 13c-d**). Based on these aligned embeddings, we transferred cell-type annotations from the CODEX slice to the Visium slice, achieving CODEX-guided spatial cell-type mapping (**Supplementary Fig. 13e, top**). The reliability of this mapping was supported by the strong spatial concordance between the predicted cell types and the expression patterns of corresponding marker genes^69,79-81^ (**Supplementary Fig. 13e, bottom**), as well as the accurate localization of germinal center (GC) cell type (follicular dendritic B cell) to histology-annotated GC zones (**Supplementary Fig. 13f)**. Collectively, these findings demonstrate STAMO’s capacity to overcome the technical limitations of existing platforms and enable a more comprehensive characterization of tissue microenvironments by integrating spatial transcriptomics and proteomics data.

### Integration of spatial DNA and RNA data for analyzing intratumor clone heterogeneity

Recently developed Slide-DNA-seq platform enables spatially resolved, genome-wide profiling of genomic alterations directly from intact tissue sections^3^. By integrating spatial DNA and RNA sequencing data, researchers can investigate both genomic mutations and their downstream transcriptomic consequences within the spatial context of tumor clones. We evaluated STAMO’s ability to integrate such datasets using a mouse liver metastasis tissue sample containing two distinct metastatic tumor clones, labeled as clone A and clone B (**Fig. 6a**). This dataset comprised two adjacent unpaired slices: one profiled by Slide-DNA-seq (copy number alteration) and the other by Slide-RNA-seq.

**Fig. 6.**
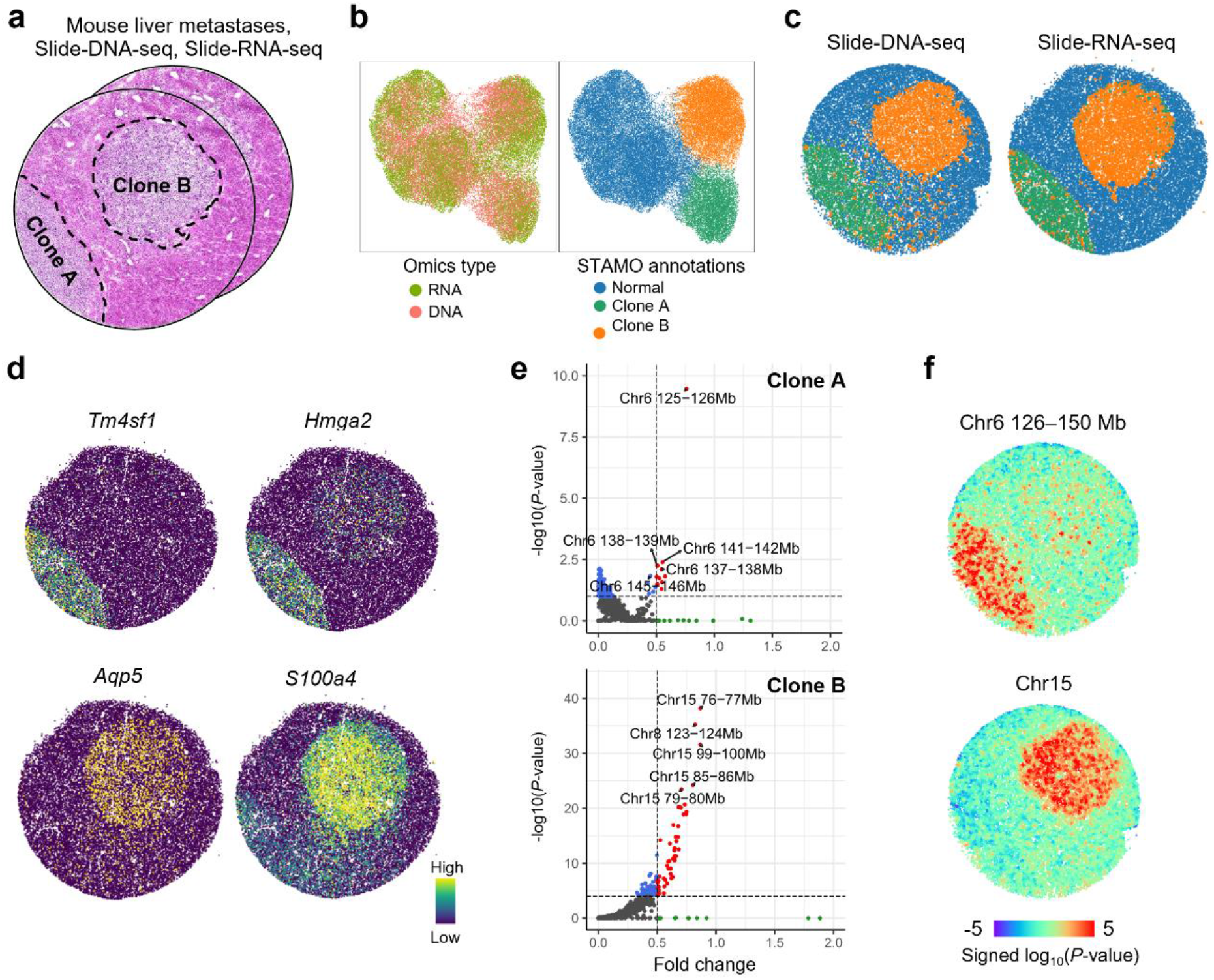
Accurate integration of spatial DNA and RNA data for analyzing intratumor clone heterogeneity. **a**. Serial slide-DNA-seq and slide-RNA-seq slice of mouse liver metastases with manual clone annotations based on H&E staining image. **b**. UMAP plots of embeddings of STAMO colored by omics type (left), and STAMO annotation (right). **c**. Visualization of aligned spatial domains across slices characterized by STAMO. **d**. Differentially expressed genes for each clone. **e**. Differential genomic copy number alterations (CNAs) for each clone. **f**. Spatial heatmaps of genomic region enrichment signed *P*-values for chromosomes 6 (126–150 Mb) and 15 (two-sided permutation test). Amplifications, red; deletions, blue.

The UMAP visualization of STAMO’s integrated embeddings revealed clear separation of the three spatial domains, while effectively mixing the two omics modalities (**Fig. 6b**). We can find that one of spatial domains identified by STAMO was visually concordant with normal tissue in H&E staining image; the other two were correspond to different metastatic clones (**Fig. 6c**). The domain annotations were further supported by the expression of established late-stage tumor markers *Tm4sf1* (JAK-STAT signaling activation) and *Hmga2* (lung metastasis signature) for clone A, *Aqp5* (loss of lineage identity) and *S100a4* (epithelial-to-mesenchymal transition) for clone B (**Fig. 6d**)^82^. Differential analysis of the copy number alteration (CNA) profiles revealed that the genomic regions with significant CNA gains for clone A and clone B was Chr 6 and Chr 15, respectively (**Fig. 6e**). We further assessed the aggregate genomic coverage in these altered regions using a two-sided Wilcoxon rank-sum test (*P*-values shown in **Fig. 6f**), providing additional evidence for the spatial distribution of these aberrations. Together, these results suggest that the spatial patterns of genomic alterations reflect the distinct evolutionary trajectories of the two tumor clones.

### STAMO enables cross-species comparison of cerebellar cortex layers and reveals conserved and primate-specific layer-enriched genes

Cross-species comparison between primates and mouse can enhance our understanding of primate-specific cerebellar functions. We applied STAMO to integrate three spatial transcriptomics cerebellar cortex slices profiled by stereo-seq from mouse, marmoset, and macaque^83^ (**Supplementary Fig. S14a**). Cross-species spatial domain alignment is usually based on homologous genes and current spatial integration approaches are mostly restricted to one-to-one homologies shared by both species, where non-one-to-one homologous genes characterizing tissue conservative features could be lost. STAMO leverage non-one-to-one (many-to-many) homologous gene mapping to construct prior feature graph. As a result, STAMO achieved the best spatial domain alignment performance when evaluating the concordance with original annotations (**Supplementary Fig. S14b-c**). Spatial visualization of the identified domains also shown high concordance with the original annotations, including large tissue structures such as granular layer (cluster 0), molecular layer (cluster 1, 2, 5, 6), white matter (cluster 4) and thin Purkinje layer (cluster 3) (**Supplementary Fig. S14d**).

We then performed a cross-species comparison of the layer-enriched genes (**Supplementary Fig. S14e**) and found that granular layer has the most primate-specific genes and molecular layer has the least conserved genes. Among the conserved genes, known layer-enriched genes such as *CBLN1*^*84*^, *PCP4*^*83*^, and *PVALB*^*84*^ were identified (**Supplementary Fig. S14f**).

The *SNCG*^*83*^ gene, encoding a synuclein family protein implicated in neurodegeneration, was preferentially enriched in Purkinje layer in primate species, whereas in mouse it demonstrated no clear laminar bias (**Supplementary Fig. S14g**). Similarly, *GRIK2* and *BTBD11* displayed enrichment in the granular and molecular layers, respectively, in primates, but lacked such layer-specific enrichment in mouse. Altogether, these species-specific, layer-enriched expression patterns point toward fundamental interspecies differences and may underlie specific functions emerging during evolution.

## Discussion

In this study, we introduce STAMO, a graph attention neural network framework, addressing the challenges in spatially aware integration of unpaired spatial multi-omics slices with partially or fully unshared features. Our results demonstrate that STAMO effectively aligns spot embeddings across different omics, preserves the biological variation, and accurately identifies consensus spatial domain. Moreover, STAMO excels in tasks of cross-omics data generation. When dealing with multi-omics data with unmatched spots, STAMO’s cross-omics generation enables the completion of unpaired multi-omics slices, thereby reducing the need for costly joint measurements. The artificially paired multi-omics data facilitates the inference of peak-gene and gene-protein links, and enables the discovery of novel gene regulatory networks that may be obscured in single-omics analyses. Additionally, with the prior feature graph where each edge is associated with a sign attribute, STAMO can integrate spatial epigenomic data with opposite regulation relationship (positive and negative). The technological advances of STAMO are summarized in **Supplementary Table 1**.

While this study primarily focuses on unpaired multi-omics integration, paired (co-profiling) datasets are becoming increasingly available. STAMO can also be applied to integrate paired multi-omics data. For paired spatial multi-omics datasets, where cell-level correspondences are known, the FGW-OT module in STAMO is not required for anchor discovery. Instead, the true anchor relationships can be directly incorporated to guide the alignment process by encouraging matched cells to be close in the shared latent space, thereby facilitating effective fusion of multi-omics information. To systematically evaluate the applicability of STAMO in this setting, we conducted benchmarking experiments on four paired mouse brain coronal spatial epigenome-transcriptome datasets, comparing against representative paired-data integration methods: SpatialGlue, SpaMosaic^23^, and MultiGATE^85^. For SpaMosaic, we adopted its vertical integration setting and included MultiGATE since it also employs a graph attention network backbone to model spatial information. As shown in **Supplementary Fig. S15**, when applied to paired datasets, STAMO achieves superior performance in spatial domain identification compared to competing methods. Visually, similar to other approaches, STAMO can clearly delineate major cortical layers. These results suggest that, although STAMO is designed for unpaired data integration, it remains competitive when applied to paired spatial multi-omics datasets.

A current limitation of STAMO is that it relies on prior feature relationships to establish cross-omics correspondences. As a result, it is not directly applicable to integrate omics such as spatial metabolomics and transcriptomics^86^, where explicit feature-level mappings (e.g., gene-metabolite relationships) are unavailable.

Looking forward, we anticipate that the increasing availability of matched histological images across spatial omics platforms may provide a promising direction for overcoming this limitation. In particular, leveraging images as omics-agnostic anchors could enable the development of new integration frameworks that do not depend on predefined feature correspondences, thereby extending STAMO to a broader range of cross-omics integration tasks.

## Methods

### Data preprocessing

The spatial multi-omics data used in this study follow a specific preprocessing pipeline for each omics. For transcriptomic data, raw gene expression counts were log-transformed and normalized by library size via the SCANPY package^87^. The top 3,000 highly variable genes were selected and used as input to the encoder. Proteomic data follows the pipeline of transcriptomic data, excluding the feature selection step. For the chromatin accessibility (peak) data, we used latent semantic indexing (LSI) to reduce the raw counts data to 50 dimensions.

### Construction of the spatial neighbor graph

For each omics slice, we construct a spot-spot spatial graph using the spatial coordinates of all spots. Pairwise similarities between spots are calculated based on their Euclidean distances. Two spots are considered neighbors if the distance between them is less than a predefined threshold *r*. This results in an undirected spatial neighbor graph represented by an adjacency matrix ***A***, where ***A***(i, j)=1 if spots i and j are spatial neighbors. The matrix ***A*** is symmetric and includes self-loops for all spots. We found that assigning 10 neighbors per spot consistently yields strong performance across datasets with varying spatial resolutions (**Supplementary Fig. S16**). In practice, the threshold *r* is determined empirically. Starting from the minimum inter-spot distance, *r* is gradually increased until each spot has approximately 10 spatial neighbors. In this study, the chosen values of *r* for different platforms are as follows: *r*=1.3 for spatial epigenome-transcriptome datasets, *r*=30 for Slide-seq, *r*=10 for Xenium and CODEX, *r* = 1.3 for Stereo-seq, and *r*=200 for 10x Visium. For convenience, we also provide an alternative function that directly specifies the number of neighbors using a k-nearest neighbors (KNN) graph, implemented as: STAMO.utils.Cal_Spatial_Net(adata, k_cutoff=N, model=‘KNN’).

### The STAMO framework

We begin by considering two omics (*K* = 2) slices, each characterized by a distinct feature matrix, denoted as 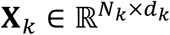, where *k* ∈ {1, 2}. Each row 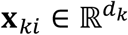 represents the feature vector of the *i*-th spot. *N*_*k*_ and *d*_*k*_ represent the number of spots and features in the slice *k*, respectively. For example, in spatial epigenome and transcriptome data integration, *d*_1_ and *d*_2_ refer to the sets of chromatin regions and genes, respectively, while for spatial transcriptomic and proteomic data integration, *d*_1_ and *d*_2_ refer to the sets of genes and proteins, respectively. As illustrated in **Fig. 1a**, the STAMO framework is composed of four modules: (1) an omics-specific variational graph attention encoder (VGAE)^88^ for spatially-aware spot embedding, (2) a prior feature graph autoencoder for feature embedding, (3) a data decoder for original omics feature reconstruction, and (4) an adversarial learning network and FGW OT module for omics alignment in the latent space. While the framework demonstrates its functionality with two omics, its modular architecture allows for straightforward extension to scenarios involving more than two omics types.

### Data encoder for spot embedding

To encode each omics data in a spatially-aware embedding space, we use a graph attention network (GAT) as the encoder of our framework^89^. It employs the graph attention mechanism to adaptively determine the influence of the neighborhood microenvironment on learning node representations and reduce the effect of the noise on the input feature data. In the following, taking the encoder for omics slice 1 as an example, we delineate the design of the VGAE module. It consists of two encoder layers. The output representation of the first encoder layer of spot *i* is defined as follows:

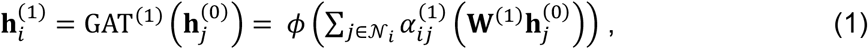

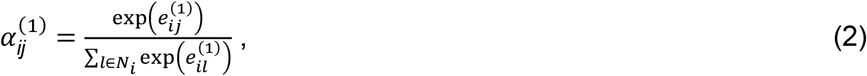

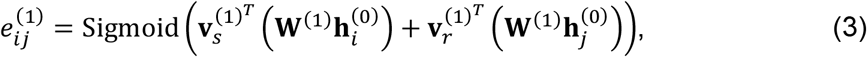

where 𝒩_*i*_ represents the neighbor set of spot *i* (including spot *i* itself), ϕ is the nonlinear activation function (LeakyReLU by default), **w**^(1)^ is the trainable weight matrix, 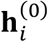 is set as the initial embeddings of the original input **x** of spot 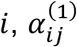 is the attention weight between spot *i* and spot *j*, measuring the relevance of the *j*-th spot on the *i*-th spot. 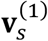 and 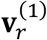 are the trainable parameters of the first layer, and Sigmoid denotes the *sigmoid* activation function.

Next, the latent representations 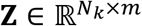 are sampled from the following normal distribution:

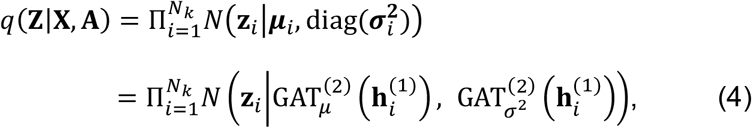

where *m* is the dimensionality of latent embedding, the mean vector **μ**_*i*_ and standard deviation vector ***σ***_*i*_ are outputs of the second encoder layer GAT^(2)^, **z**_*i*_ ∈ ℝ^*m*^ corresponds to the latent representation of the *i* -th spot. Sampling directly from this distribution during model training introduces stochasticity, which hinders the backpropagation process because gradients cannot flow through random sampling operations. To overcome this, **z**_*i*_ can be obtained through reparameterization^90^ by **z**_*i*_ = **μ**_*i*_ + ***σ***_*i*_ ⨀ **ϵ**, where the noise variable **ϵ** ∼ *N*(**0, I**).

The loss function of the VGAE module consists of two parts. The first part is the spatial graph regularization loss, which makes the closeness of embedding points similar to the spatial proximity of spots. It uses the binary cross-entropy loss to measure the difference between the input adjacency matrix and the reconstructed adjacency matrix. The reconstructed adjacency matrix is generated by an inner product between latent variables as follows:

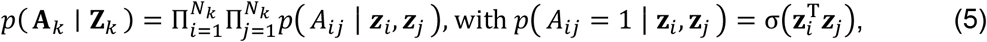

where *σ*(·) is the logistic sigmoid function. The second part (latent distribution regularization) is the Kullback-Leibler (KL) divergence between *q* (**Z**|**X, A**) and the Gaussian prior *p*(**Z**) = Π_*i*_ *p*(***z***_*i*_) = Π_*i*_*N*(***z***_*i*_|0, **I**). The formulation of the two parts can be written in the following form:

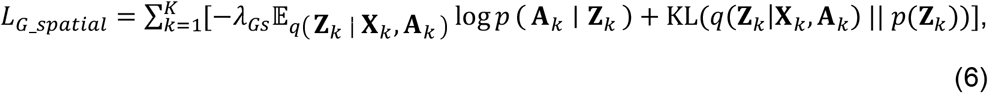

where *λ*_*Gs*_ is the parameter to control the spatial regularization strength.

### Prior feature graph construction

To enable semantically consistent spot embeddings from different omics encoders, STAMO adopts the feature graph-linked embedding learning framework from GLUE^12^. Specifically, the prior feature graph is denoted as 𝒢 = (𝒱, ℰ), where 𝒱 = {𝒱_1_, …, 𝒱_*K*_} represents the set of omics features (e.g., genes for RNA-seq, peaks for CUT&Tag /ATAC-seq, and genome bins for DNA-seq) and ℰ represents the set of edges capturing regulatory relationships between features from different omics layers.

In STAMO, cross-omics feature relationships are systematically established based on genomic proximity. Specifically, we obtain genomic annotation data from the GENCODE^91^ database and extract genomic coordinates for features from each omics modality. Edges are then defined between features whose genomic coordinates overlap or fall within a specified distance. For example, ATAC peaks are linked to genes if they overlap with gene bodies or promoter regions; protein features are mapped to their corresponding coding genes via established protein-gene relationships; and DNA features are segmented into genomic bins that are linked to overlapping RNA genes.

This graph construction strategy naturally generalizes to integration scenarios involving more than two omics modalities, as edges are consistently defined based on genomic distance or overlap. In such cases, transcriptomic genes can serve as intermediate anchors to connect features across modalities (e.g., linking chromatin accessibility and histone modification features), enabling flexible graph construction for diverse omics combinations. For cross-species integration, we construct feature correspondences based on gene homology using the biomaRt^92^ package. Importantly, the retrieved mappings include one-to-one, one-to-many, and many-to-many relationships. Rather than enforcing one-to-one constraints, we retain all homologous gene pairs and encode each pair as an edge in the prior graph. This design allows genes to connect to multiple homologs across species, thereby capturing complex evolutionary relationships and preserving conserved biological signals for downstream spatial alignment.

To model the directionality of the regulatory effect, each edge is associated with a sign attribute *s*_*i,j*_ ∈ {−1, 1}. For instance, ATAC peaks are generally associated with the activation of gene expression (denoted as *s*_*i,j*_ = 1), whereas histone modification H3K27me3 or DNA methylation in the gene promoter are typically considered repressive to gene expression (denoted as *s*_*i,j*_ = −1). In addition to the connections between features, self-loops are also added for numerical stability, with *s*_*i,i*_ = 1, ∀*i* ∈ 𝒱.

We conducted benchmarking experiments to systematically evaluate the robustness of STAMO under noisy, incomplete, and partially incorrect prior feature graphs. STAMO exhibited only a modest decline in performance provided that the proportion of noisy edges remained below 80% of the true edges, or that fewer than 80% of edges were deleted or replaced, demonstrating its strong robustness (**Supplementary Fig. S17**).

### Prior feature graph autoencoder

The feature embeddings **F** ∈ ℝ^*m*×|𝒱|^ are initialized with zero vectors and updated by a graph convolutional network (GCN):

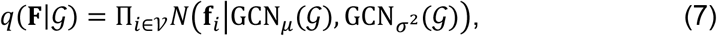

where **f**_*i*_ ∈ ℝ^*m*^. Similar to the spatial neighbor graph reconstruction, the feature graph likelihood is defined as: 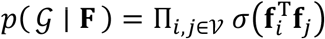. The loss function for feature graph reconstruction can be written in the following form:

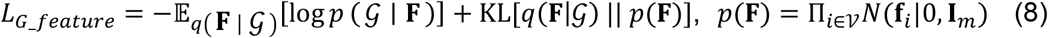

### Data decoder for reconstruction

The data likelihoods *p*(**x**_*k*_ ∣ **z, F**_*k*_), serving as data decoders, are modeled based on the inner product between the spot embedding **z** and feature embedding 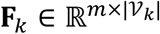. The specific form of the likelihood function is determined by the statistical characteristics of the omics data. For instance, for count-based omics data such as spRNA-seq and spATAC-seq data, the negative binomial (NB) distribution^16^ is employed:

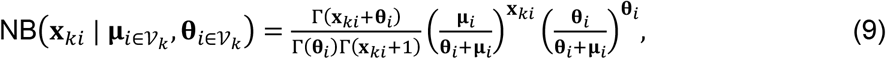

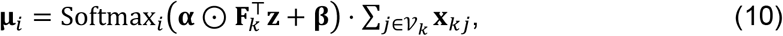

where 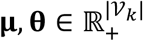 are the mean and dispersion of the NB distribution,respectively. The vector 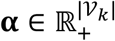 and 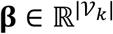 indicate scaling and bias factors, respectively, while ⊙ denotes the Hadamard product, Softmax_*i*_ indicates the *i*th dimension of the softmax output and 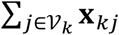 represents the total count within a given spot. Applying the softmax function followed by multiplication with the total count ensures that the reconstructed data preserves the original library size. The negative log likelihood of the NB distribution can be used as the reconstruction loss function of the original data **x**_*k*_, and can be written in the following form:

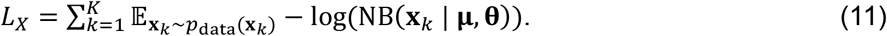

### Adversarial learning-based alignment

To align the latent embeddings generated from different omics modalities, we adopt an adversarial learning strategy, which has been proven effective in prior studies^93-95^. A discriminator network D is introduced to infer the omics types of spots based on their embedding **z**. The discriminator D is trained to minimize the following multi-class cross-entropy loss:

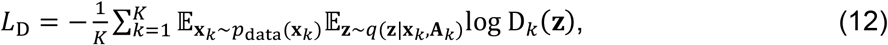

where D_*k*_ denotes the *k* th dimension of the discriminator’s output, corresponding to the predicted omics label. The discriminator and encoder are trained in an adversarial manner: the discriminator is optimized to minimize *L*_D_, while the encoder is trained to maximize it. This adversarial interplay forces the encoder to produce omics-invariant embeddings, thereby promoting alignment of spot embeddings from different omics slices.

### FGW OT-based anchor identification

Since adversarial learning can only align the embedding in a global manner, we further adopt the Fused Gromov-Wasserstein optimal transport^96^ to identify anchors across the omics slice and then use these guide anchors to achieve the local alignment. Since different omics slices do not share the original feature space, we define the transport cost between different slices in the common latent space produced by the encoder. The cost between spot *i* in slice 1 and spot *j* in slice 2 is defined as the Euclidean distance of latent vectors **z**_1*i*_ and **z**_2*j*_ as follows: *c* (**z**_1*i*_, **z**_2*j*_) = ||**z**_1*i*_ − **z**_2*j*_||. We then compute the following transport plan:

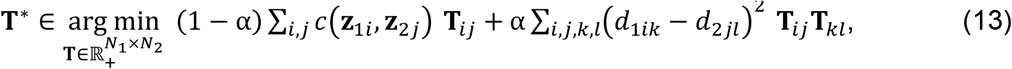

where the first term quantifies the embedding distance between two slices and the second term quantifies the structural correspondence between the spatial graphs of two slices. Here, **T**_*ij*_ represents the mapping/alignment probability between spot *i* in slice 1 and spot *j* in slice 2, *d*_1*ik*_ represents the spatial coordinate distance between spot *i* and *k* on the slice 1, and *α* balances the importance between the expression profile and the spatial graph structure.

We compared FGW-OT with two prior optimal transport-based alignment approaches developed for single-cell multi-omics integration: Wasserstein OT (W-OT), as used in UniPort^97^, and Gromov-Wasserstein OT (GW-OT), as used in SCOT^25^. W-OT measures only expression distances between datasets, whereas GW-OT focuses solely on structural correspondence and preserves local geometry. FGW-OT integrates both aspects, and we introduce a weighting parameter *α* to balance their contributions. Using spatial domain matching accuracy as the evaluation metric, our results (**Supplementary Fig. 18**) demonstrate that FGW-OT significantly improves anchor identification accuracy compared to OT methods that consider only expression or structure. With the optimal spot-spot mapping matrix **T**^*^, we identify for each spot its cross-slice anchor with the highest matching probability and remove any duplicated anchors. These anchor spot pairs are then used to guide the alignment process by encouraging them to be close to each other in the common latent space. This is achieved by adding the following mean squared error loss:

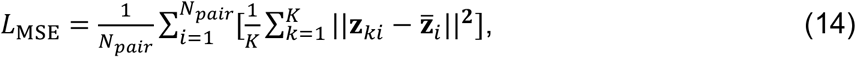

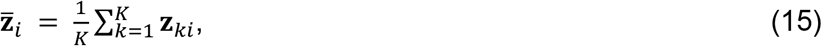

where *N*_*pair*_ is the size of anchor pairs.

In summary, the final objective function is written as:

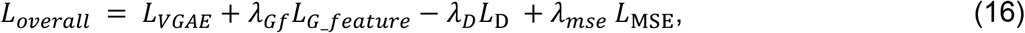

where *λ*_*Gf*_, *λ*_*D*_, and *λ*_*mse*_ are weight parameters and *L*_*VGAE*_ = *L*_*X*_ + *L*_*G*_*spatial*_. STAMO is trained in a two-stage workflow. In the first stage, we train the model to generate preliminary aligned embeddings by setting *λ*_*mse*_ = 0. In the second stage, anchors are identified based on the embeddings and then used to refine the alignment process. The pseudocode of the training procedure is summarized in the **Supplementary Note**.

### Cross-omics data generation

Once the training is finished, the network compositions G2(E1(**X**_1_)) and G1(E2(**X**_2_)) can be used to generate unmeasured features across omics, where E1 and E2 are graph encoders for omics 1 and 2, and decoders G1 and G2 are employed to generate spot features from the latent embeddings. In this way, the model transfers information across omics via the aligned latent space while preserving spatial context through the graph-based autoencoders.

### Implementation details of STAMO

For each omics, the encoder consists of two layers with node dimensions set as 256 and 50, respectively. We use the PyTorch Geometric library^98^ to implement the graph attention layer. We use the RMSprop optimizer with a learning rate of 0.002 and a minibatch size of 128 to train it. The weight factors *λ*_*Gs*_, *λ*_*Gf*_, *λ*_*D*_, and *λ*_*mse*_ are set as [0.001, 0.02, 0.05, 0.02] for all datasets. We set the trade-off parameter *α* = 0.5 by default. STAMO demonstrates strong robustness to hyperparameter variation, with its default configuration achieving performance close to optimal across all evaluated datasets (**Supplementary Fig. 19**).

To overcome the quadratic time and memory complexity of FGW-OT, we employed the recently proposed low-rank FGW formulation, which constrains the transport coupling matrix to be low-rank and enables linear time and memory complexity. We implement this approach using Optimal Transport Tools (OTT)^99^, a scalable JAX-based library that supports just-in-time (JIT) compilation and efficient on-the-fly evaluation of the cost function.

We first evaluated whether the low-rank approximation compromises performance. Specifically, we compared anchor identification accuracy between full-rank and low-rank FGW under different rank settings (10, 100, and 1,000; **Supplementary Fig. 20a**). The results show that a rank of 1,000 achieves accuracy comparable to the full-rank formulation. Therefore, for large-scale datasets, we use rank = 1,000 as it provides the best trade-off between accuracy and efficiency among the tested configurations. For smaller datasets (fewer than 10K cells), STAMO is executed in full-rank mode to retain maximal precision.

We further conducted runtime and memory benchmarks on the human lymph node dataset, the largest dataset used in this study, which contains a total of 566K cells, including 377,897 cells from a 10x Xenium slice and 188,450 cells from a CODEX slice. As shown in **Supplementary Fig. 20b**, the runtime and memory usage of low-rank FGW scale approximately linearly with dataset size, whereas full-rank FGW exhibits quadratic scaling and is unable to complete the integration beyond 250K cells. Notably, when applied to the full dataset (566K cells), STAMO solves the FGW-OT problem in approximately 0.43 hours while using only 10.2 GB of system memory.

### Clustering

For datasets lacking ground-truth annotations, we employ the Leiden algorithm implemented in the SCANPY package with the default settings for clustering. To capture known biological structures, we evaluate clustering outcomes across a range of cluster numbers. For datasets with ground-truth labels, we employ a binary search strategy to tune the resolution parameter of the Leiden algorithm, aiming to match the number of spatial clusters with the number of ground-truth annotations. The procedure starts by setting a minimum and maximum resolution. In each iteration, clustering is performed using the midpoint of the current resolution range. If the resulting number of clusters exceeds the ground truth, the maximum resolution is updated; otherwise, the minimum is updated. This process continues until the number of clusters matches the ground truth.

### Evaluation metrics

The quality of the embeddings provided by benchmarking methods are assessed in three aspects: Spatial domain identification is assessed using the adjusted rand index (ARI) and Moran’s *I* metrics^100^, batch correction is assessed using mean average precision (MAP), average silhouette width (ASW), graph connectivity (GC), seurat alignment score (SAS), and modality alignment is assessed using Fraction of Samples Closer than True Match (FOSCTTM)^12^.

*Moran’s I*: Moran’s *I* score assesses whether the spatial pattern of spatial domain labels is clustered, dispersed, or random, which helps in understanding the spatial continuity within identified regions. Given *N* spots with their neighboring weight graph 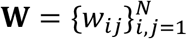 and associated spatial domain labels 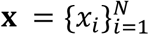 with their mean 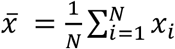, the Moran’s *I* statistic is defined as:

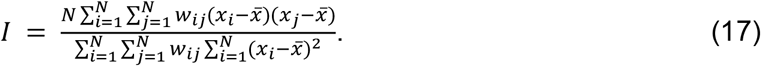

Moran’s *I* ranges from -1 to 1, where higher values indicate stronger spatial autocorrelation and a smoother spatial pattern.

#### Mean average precision (MAP)

MAP is used to evaluate the spatial domain resolution. Let *y*^(*i*)^ denote the spatial domain label of the *i* th spot, and let 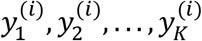 represent the spatial domain labels of its *K* nearest neighbors, ranked by proximity. The MAP score is defined as:

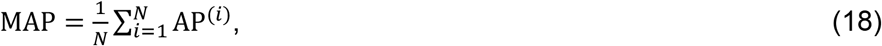

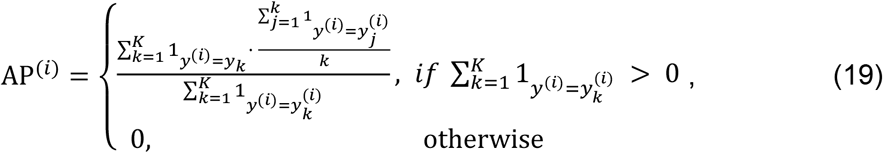

where 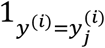 is an indicator function that returns 1 if 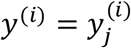 and 0 otherwise. For each spot, average precision (AP) measures the average proportion of label-consistent neighbors among the top-*K* ranked neighbors. MAP is then computed as the average AP over all spots. In our experiments, we set *K* to 1% of the total number of spots in each slice. MAP ranges from 0 to 1, with higher values indicating better spatial domain resolution.

#### Average silhouette width (ASW) for spatial domain label

Domain label ASW is also used to assess spatial domain resolution. It is computed as:

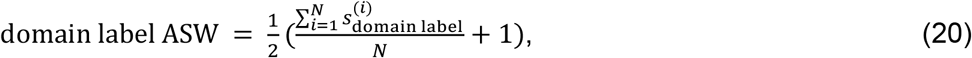

where 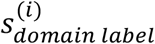 denotes the spatial domain label silhouette width for the *i*th spot, and *N* is the total number of spots. Domain label ASW ranges from 0 to 1, with higher values indicating better spatial domain resolution.

#### Average silhouette width (ASW) for omics type label

Omics type ASW is used to evaluate the degree of mixing across omics types. It is computed as:

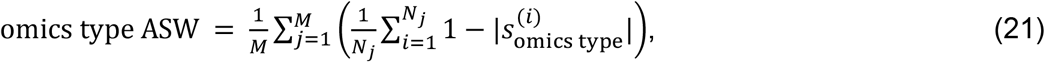

where 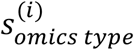 denotes the omics type silhouette width for the *i*th spot, *N* is the number of spots in the spatial domain *j*, and *M* is the total number of spatial domains. Omics type ASW ranges from 0 to 1, with higher values indicating better mixing.

#### Graph connectivity (GC)

GC is used to evaluate the degree of mixing across omics types within each spatial domain. It is defined as:

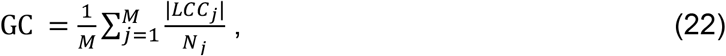

where *LCC*_*j*_ denotes the number of spots in the largest connected component of the *k*-nearest neighbor graph (*K* = 15) within the spatial domain *j, N*_*j*_ is the total number of spots in domain *j* and *M* is the total number of spatial domains. GC ranges from 0 to 1, with higher values indicating better mixing.

#### Seurat alignment score (SAS)

SAS quantifies the degree of omics type mixing by measuring the composition of each spot’s neighborhood. It is defined as:

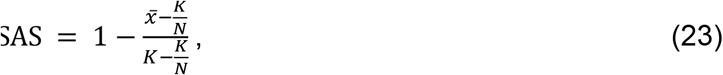

where 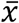 denotes the average number of spots from the same omics type among the *K* -nearest neighbors (different omics types are subsampled to match the number of spots in the smallest type to ensure balanced comparison), and *N* is the number of omics types. We set *K* to 1% of the subsampled spot number. SAS ranges from 0 to 1, with higher values indicating better mixing.

#### Biology conservation and Batch mixing

Based on the metrics described above, the biological conservation score can be computed as follows^12^:

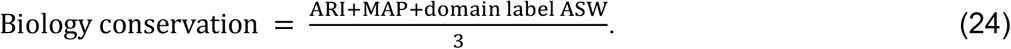

For embryonic mouse brain datasets, the batch label includes different developmental stages and omics types. Thus, we define the batch mixing score as follows:

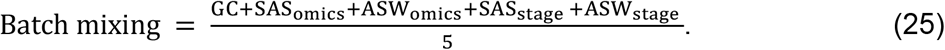

#### FOSCTTM

FOSCTTM evaluates the accuracy of single-spot-level alignment of different omics data with ground-truth pairing information. This metric has been previously applied to benchmark diagonal integration methods on paired multi-omics data^25^. Let **x**_*i*_ and **y**_*i*_ denote the embeddings of the *i*th spot from the first and second slices, respectively. FOSCTTM evaluates, for each spot in one slice, how many spots in the other slice are incorrectly ranked closer than their true counterparts. It is defined as:

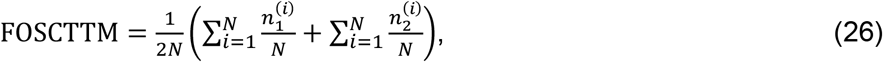

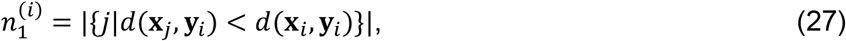

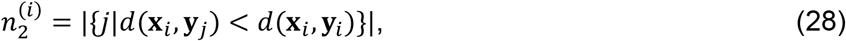

where *d* denotes the Euclidean distance. FOSCTTM ranges from 0 to 1, with lower values indicating more accurate alignment between matched spots.

### Benchmarking integration methods

As highlighted among the top-performing methods for single-cell and spatial multi-omics data integration in recent reviews^101,102^, the following representative integration methods were selected for benchmarking: Seurat, scVI, STAligner, CellCharter, and GLUE. Additionally, we produced uncorrected embeddings of the combined raw input data using SCANPY to demonstrate batch effects, where principal component analysis (PCA) was used to perform dimensionality reduction without applying any further integration algorithm. For all methods, default parameter settings were used.

To integrate spRNA and spATAC data with Seurat, scVI, CellCharter, and STAligner, requiring shared features as input, we first used Signac^103^ to transform spATAC data into a gene activity score matrix. We then extracted the set of common genes shared between the gene activity scores and the gene expression matrix as input. spCUT&Tag data were preprocessed similarly.

### Preprocessing slide-DNA-seq data

DNA features were segmented into non-overlapping 1-Mb genome bins, and copy number values were aggregated accordingly. To construct the prior feature graph, each genomic bin was linked to RNA genes that overlapped with it, either within the gene body or proximal promoter regions. The resulting bin-level copy number values were then preprocessed in the same manner as ATAC data and subsequently used as input to the model.

### Differential expression analysis

We used the scanpy.tl.rank_genes_groups() function from SCANPY to identify DEGs of the spatial domain. Genes with log2foldchange > 1 and adjusted *P* < 10^−5^ were selected as marker genes for each spatial domain. To analyze the correlation between CSS/GAS and gene expression, we divided the genes into four sections based on their log2(fold change). Specifically, genes with positive log2(fold change) in both RNA and ATAC data were assigned to quadrant I; the remaining genes were categorized into quadrants II, III, and IV accordingly based on the signs of their respective fold changes.

### Deviation score

The deviation score is designed to estimate transcription factor activity at the single-cell level from sparse chromatin accessibility data. In our analysis, the deviation score was calculated using the addDeviationsMatrix function implemented in ArchR^104^.

## Data availability

The spatial epigenome-transcriptome (ATAC-RNA and CUT&Tag-RNA) dataset of mouse postnatal day 22 brain can be accessed from Gene Expression Omnibus with accession code GSE205055. The MISAR-seq (ATAC-RNA) dataset of the mouse embryo can be downloaded from the National Genomics Data Center with accession number OEP003285. The slide-DNA-seq and slide-RNA-seq dataset of mouse liver metastases is available at the Broad Institute Single Cell Portal (https://singlecell.broadinstitute.org/single_cell/study/SCP1278). The 10x Xenium and Visium spatial transcriptomics slices of adult human lymph node are downloaded from the 10x Genomics website https://www.10xgenomics.com/datasets/human-lymph-node-preview-data-xenium-human-multi-tissue-and-cancer-panel-1-standard and https://support.10xgenomics.com/spatial-gene-expression/datasets/1.1.0/V1_Human_Lymph_Node, respectively. The CODEX spatial proteomics slice of an adult human lymph node is downloaded from the HuBMAP repository with Dataset ID HBM754.WKLP.262. The spatial transcriptomics (Stereo-seq) slices of macaques, marmosets, and mice cerebellar cortex can be downloaded from http://db.cngb.org/stomics/cbmsta. A summary of these datasets can be found in **Supplementary Table 2**.

## Code availability

An open-source Python implementation of the STAMO package and tutorials for reproducing the results are available at https://github.com/zhanglabtools/STAMO.

## Author contributions

S.Z. and L.C. conceived and supervised the project. X.Z. developed and implemented the STAMO algorithm. X.Z., K.D., J.X., L.C., and S.Z. validated the methods and wrote the paper. All authors read and approved the final paper.

## Acknowledgements

This work has been supported by National Key R&D Program of China (no. 2025YFF1207900 to S.Z. and L.C.), Natural Science Foundation of China (nos. 32341013, 12326614 to S.Z., and T2350003, T2542018, 12131020, 42450084, 42450135, 12326614, 12426310 to L.C.), the CAS Project for Young Scientists in Basic Research (no. YSBR-034 to S.Z.), the Tianfu Jincheng Laboratory (no. TFJCPI20260001 to L.C.), Zhejiang Province Vanguard Goose-Leading Initiative (no. 2025C01114 to L.C. and S.Z.), Hangzhou Institute for advanced study of UCAS (no. 2024HIAS-P004 to L.C.), Shenzhen Medical Research Fund (no. E250200621 to L.C.), and JST Moonshot R&D (no. JPMJMS2021 to L.C.).

