## Supplemental Text and Figures for "Deciphering cross-omics complexity of tissues via diagonal integration of unpaired spatial multi-omics data"

#### Supplementary Notes

The pseudocode for STAMO is summarized in **Algorithm 1**.

**Algorithm 1** STAMO algorithm

**Input:** Slice 1 of omics type 1 and Slice 2 of omics type 2 with feature matrix $\mathbf{X}_k = [\mathbf{x}_{k1}, \ldots, \mathbf{x}_{kN_k}] \in R^{N_k \times d_k}, k = 1, 2$, spatial adjacency matrix $\mathbf{A}_k$ computed by spatial coordinates, and prior feature graph $\mathcal{G}$.

**Output:** aligned latent vectors $\mathbf{z}$.

**STAMO performs the following steps:**

**1.** Initialize graph autoencoder parameters.

**2.** For $epoch = 1 \text{ to } P$ do the following:

**2.1.** Compute the hidden representation $\mathbf{Z}_k$ of spots by the data encoder and the feature embedding $\mathbf{F}_k$ by the feature encoder.

**2.2.** Reconstruct $\mathbf{A}_k$ by $\mathbf{Z}_k$ and $\mathbf{X}_k$ by the inner product between $\mathbf{Z}_k$ and $\mathbf{F}_k$ with the data decoder.

**2.3.** Compute total loss $L_{overall} = L_{VGAE} + \lambda_{Gf} L_{G\_feature} - \lambda_D L_{\mathrm{D}}$ and update parameters with gradient descent.

**3.** Based on the aligned latent vectors $\mathbf{z}$, compute the optimal cross-omics mapping matrix $\mathbf{T}^*$ by FGW OT and identify spot anchors. Repeat step 2 with the new total loss $L_{overall} = L_{VGAE} + \lambda_{Gf} L_{G\_feature} - \lambda_D L_{\mathrm{D}} + \lambda_{mse} L_{\mathrm{MSE}}$.

### Supplementary Figures

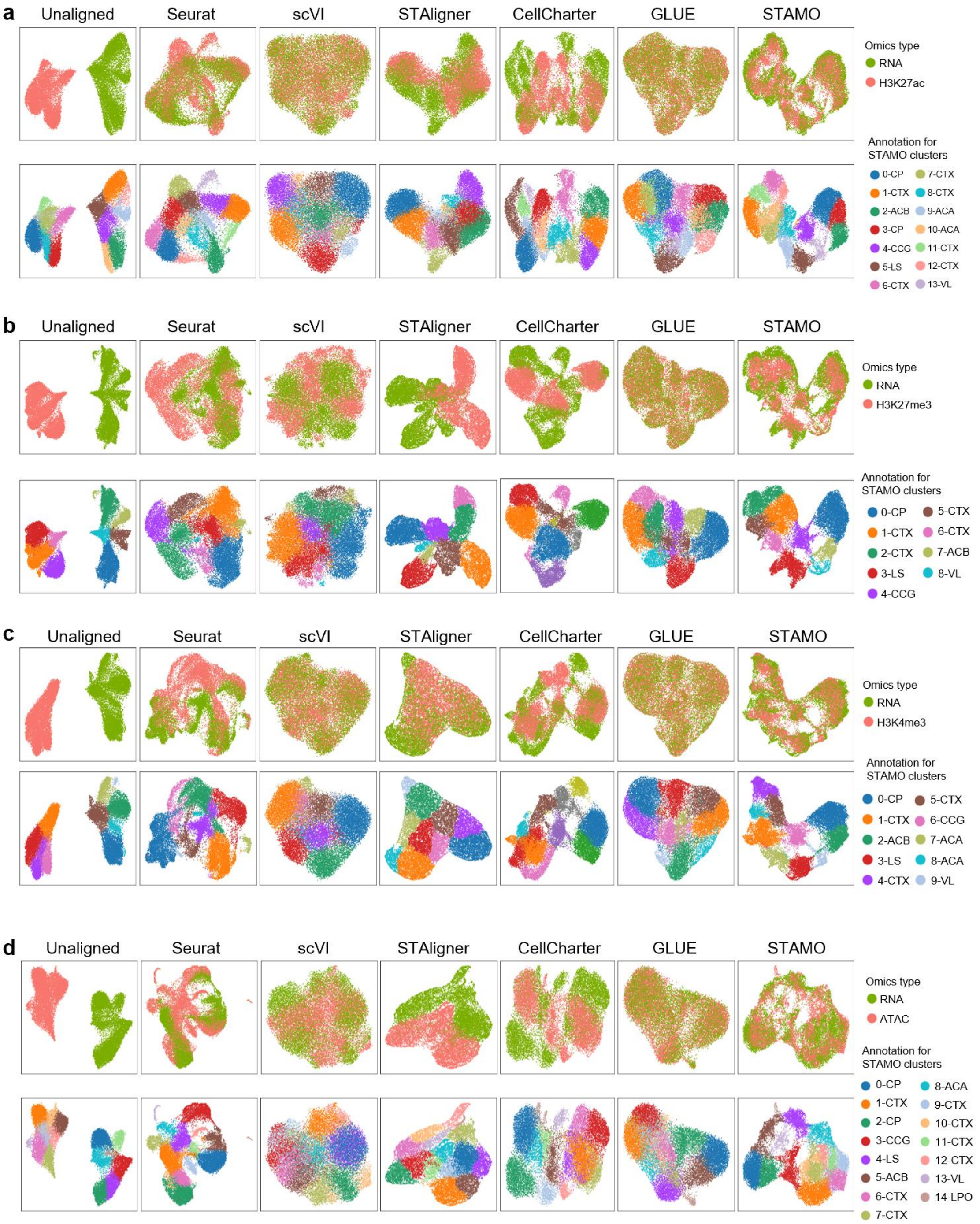


**Supplementary Fig. S1.** UMAP plots of the embeddings from all methods. Spots are colored by omics type (top) and colored by cluster (bottom). Integration results of spH3K27ac and spRNA dataset (**a**), spH3K27me3 and spRNA dataset (**b**), spH3K4me3 and spRNA dataset (**c**), spATAC and spRNA dataset (**d**), are shown.

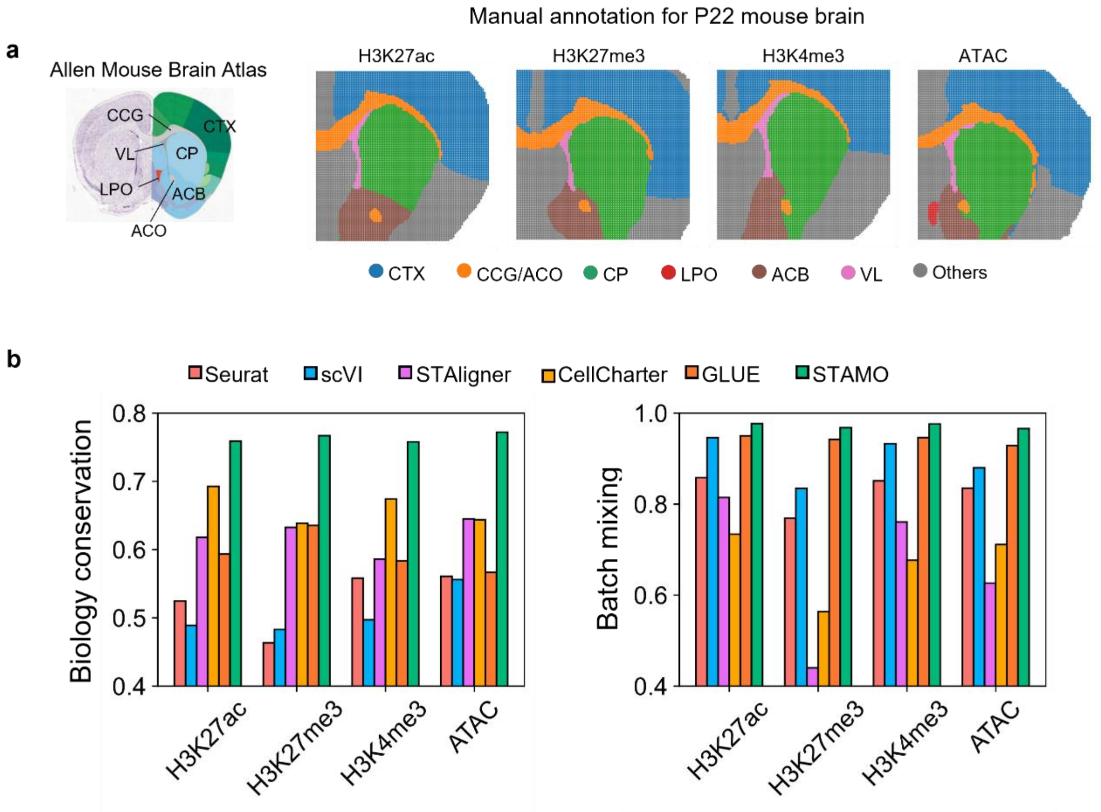


**Supplementary Fig. S2. a.** Annotation of mouse brain coronal section from the Allen Brain Atlas (left) and manual annotations of the four mouse brain coronal sections (right). **b.** Comparison of STAMO with competing methods using biology conservation and batch mixing metrics.

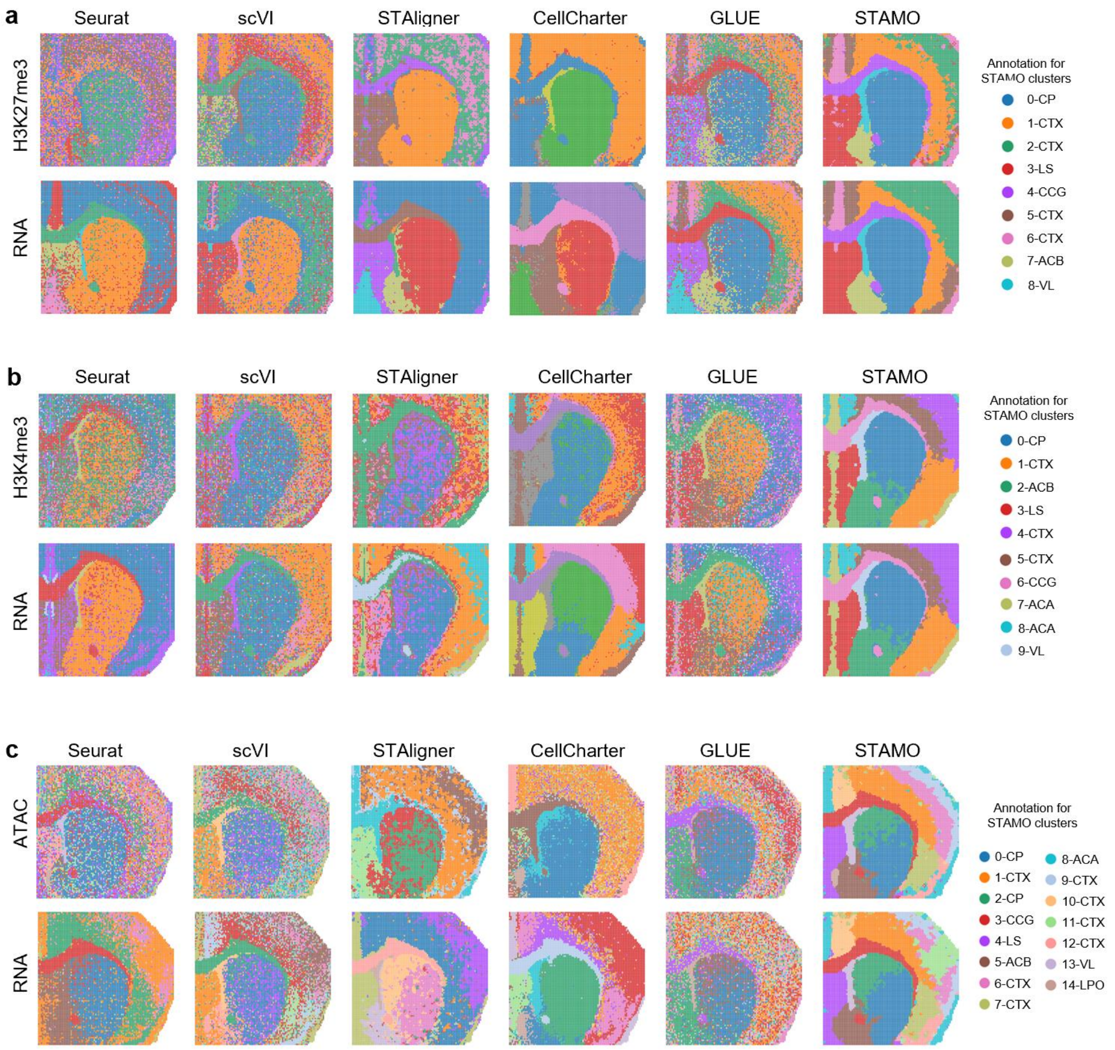


**Supplementary Fig. S3.** Spatial domains characterized by different integration methods on spH3K27me3 and spRNA dataset (**a**), spH3K4me3 and spRNA dataset (**b**), spATAC and spRNA dataset (**c**).

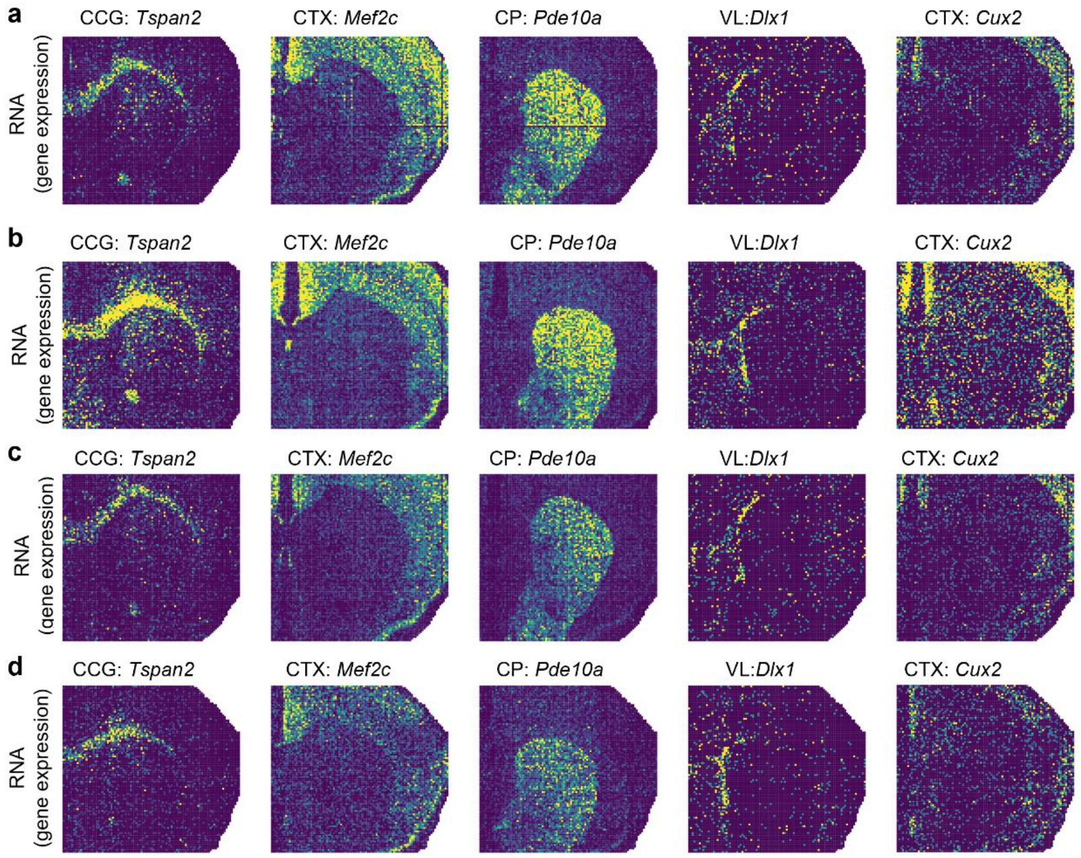


**Supplementary Fig. S4.** Spatial heatmap of gene expression of marker genes for annotated spatial domains on H3K27ac dataset (**a**), H3K27me3 dataset (**b**), H3K4me3 dataset (**c**), ATAC dataset (**d**).

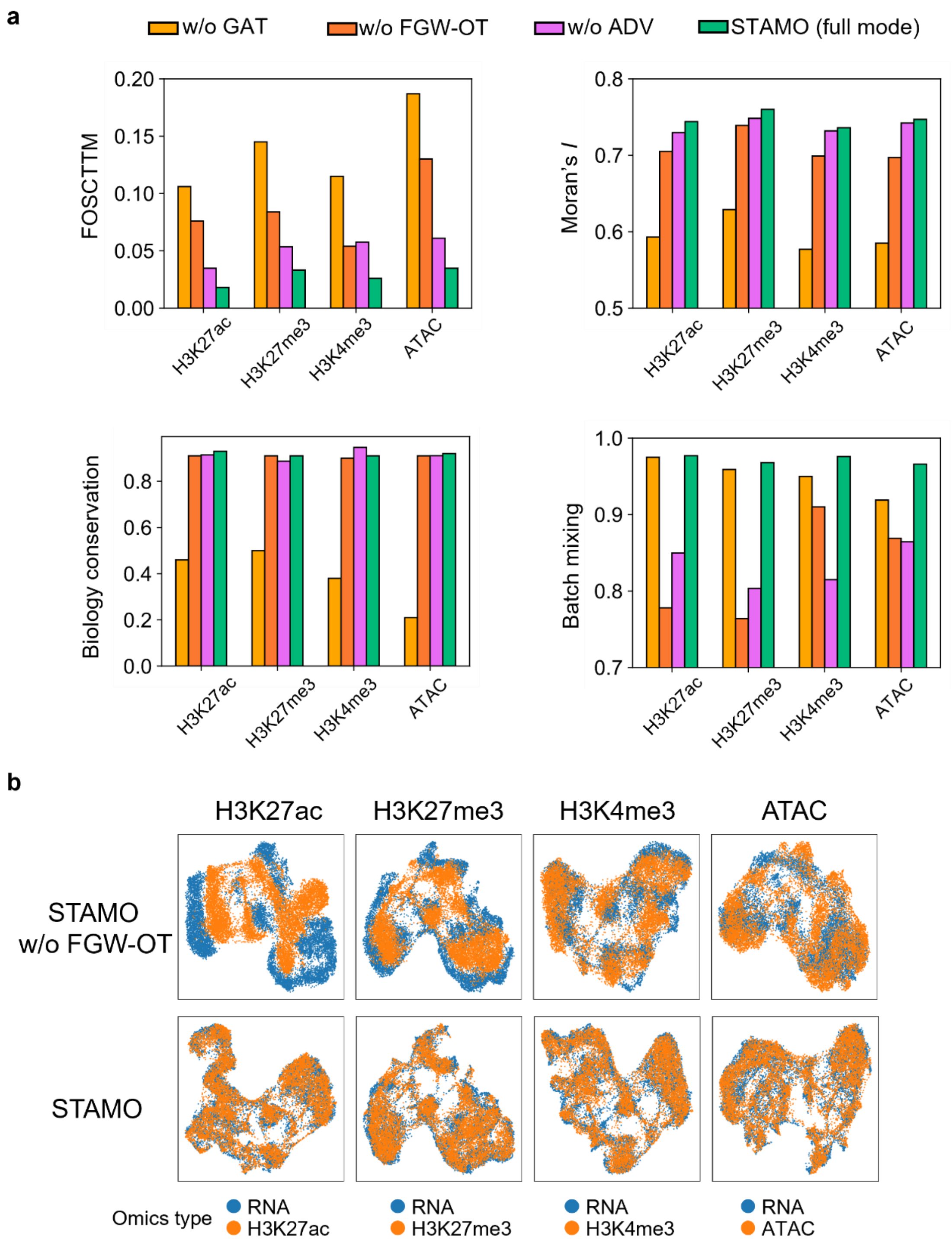


**Supplementary Fig. S5.** Ablation study evaluating the contribution of omics-specific variational graph encoder (GAT), FGW-OT, and adversarial learning discriminator (ADV) in STAMO. **a.** Integration performance of different ablation settings by removing GAT, FGW-OT, and ADV. **b.** UMAP plots of STAMO (w/o FGW-OT) and STAMO (full mode) colored by omics type.

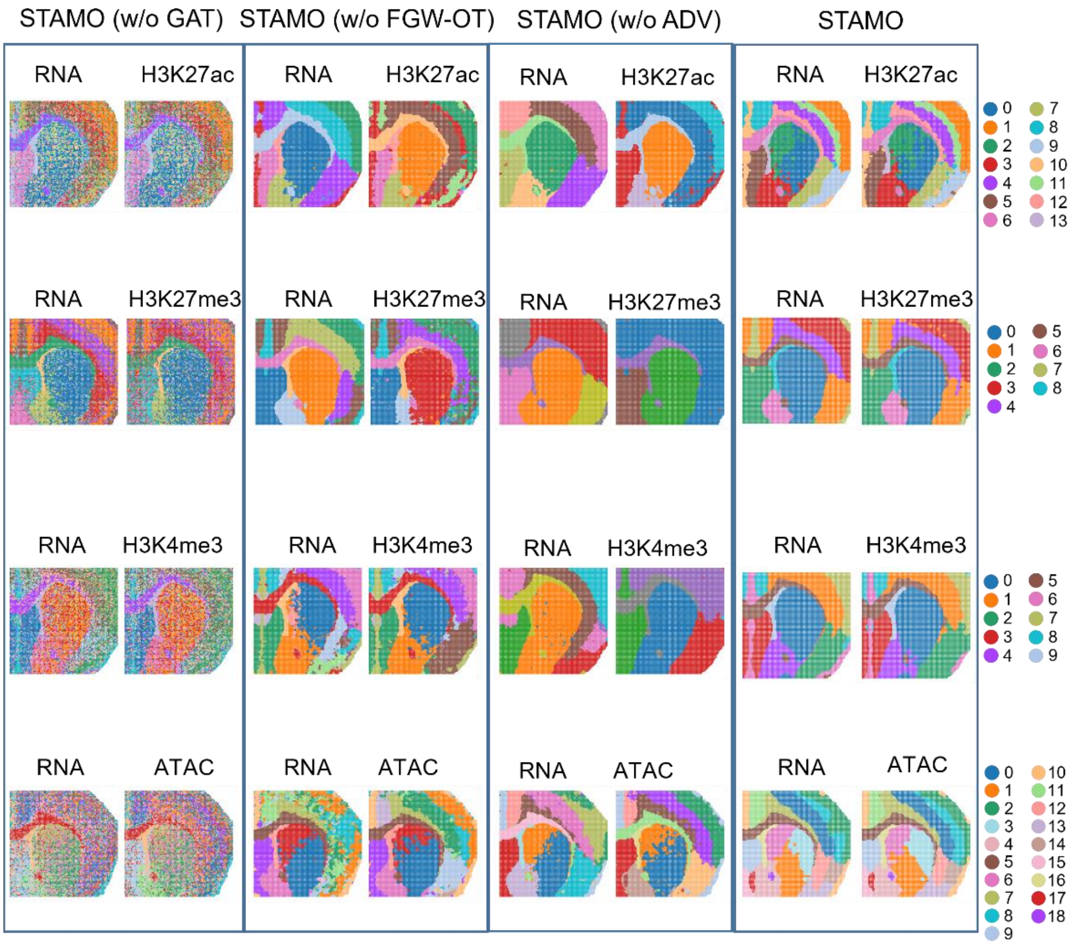


**Supplementary Fig. S6.** Spatial domains characterized by STAMO with different ablation settings (removing GAT, FGW-OT, and ADV).

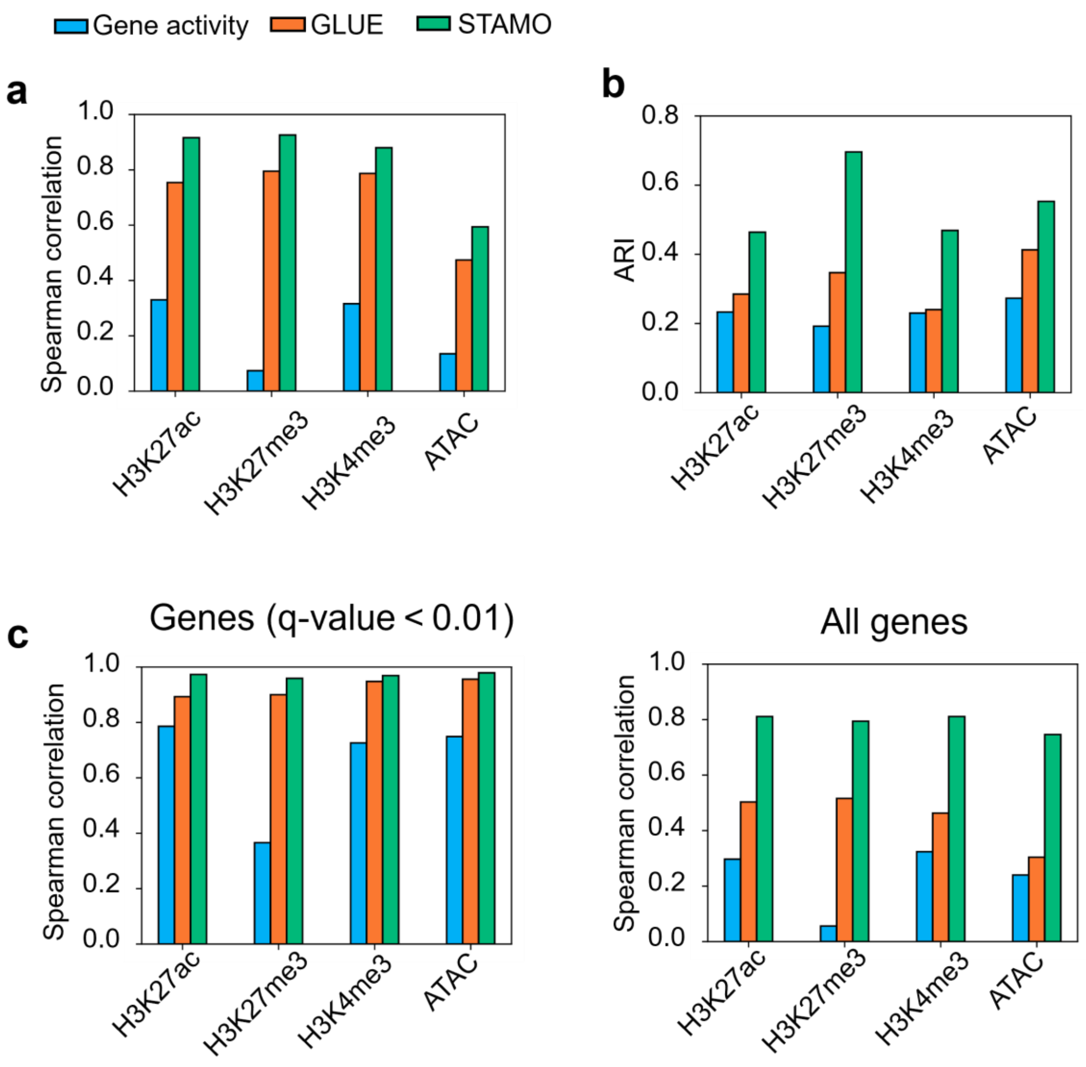


**Supplementary Fig. S7. Evaluation of cross-omics generation performance. a.** Preservation of gene-gene correlation structure, quantified by the Spearman correlation between gene-gene correlation matrices derived from generated and measured gene expression data. **b.** Spatial domain identification performance using generated data. **c.** Consistency of differential analysis between generated and measured expression, assessed by log fold-change (logFC) for genes with high confidence (q-value < 0.01, left) and for all genes (right).

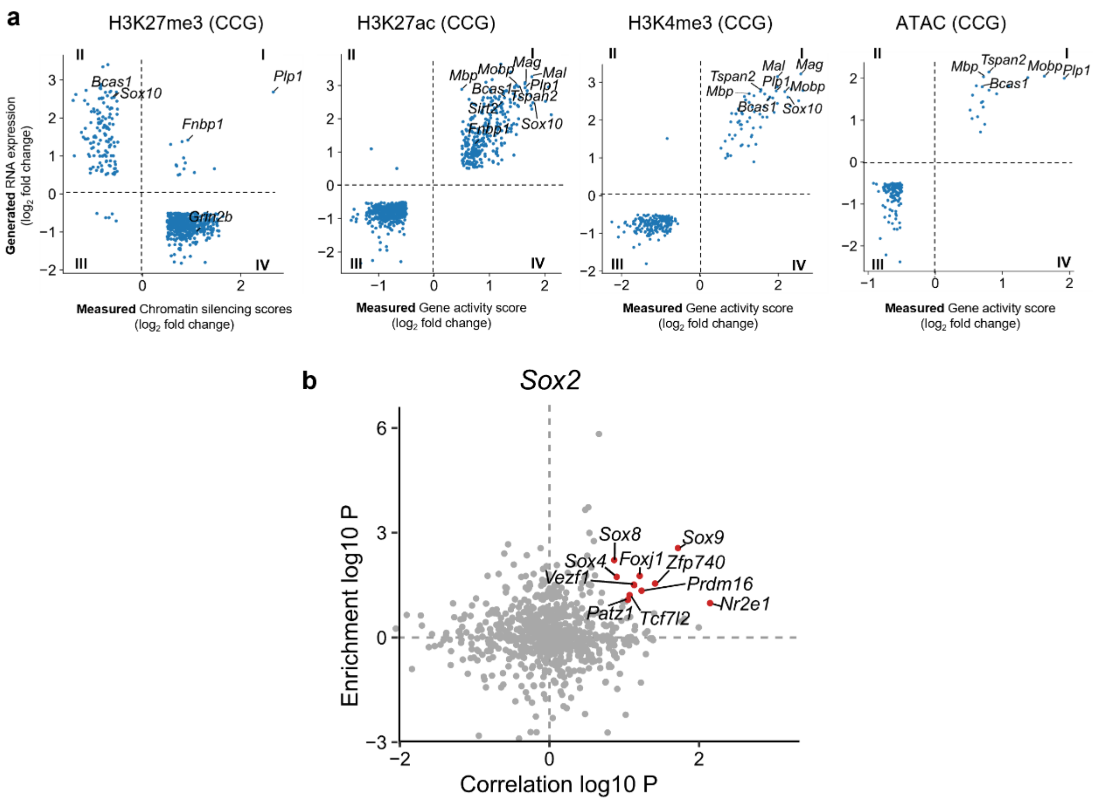


**Supplementary Fig. S8. a.** Correlation between generated RNA gene expression and measured gene activity score, chromatin silencing scores in the CCG region. **b.** Candidate TF regulators of *Sox2*. Highlighted points are TFs with an absolute regulation score ≥ 0.5, with all other TFs shown in gray.

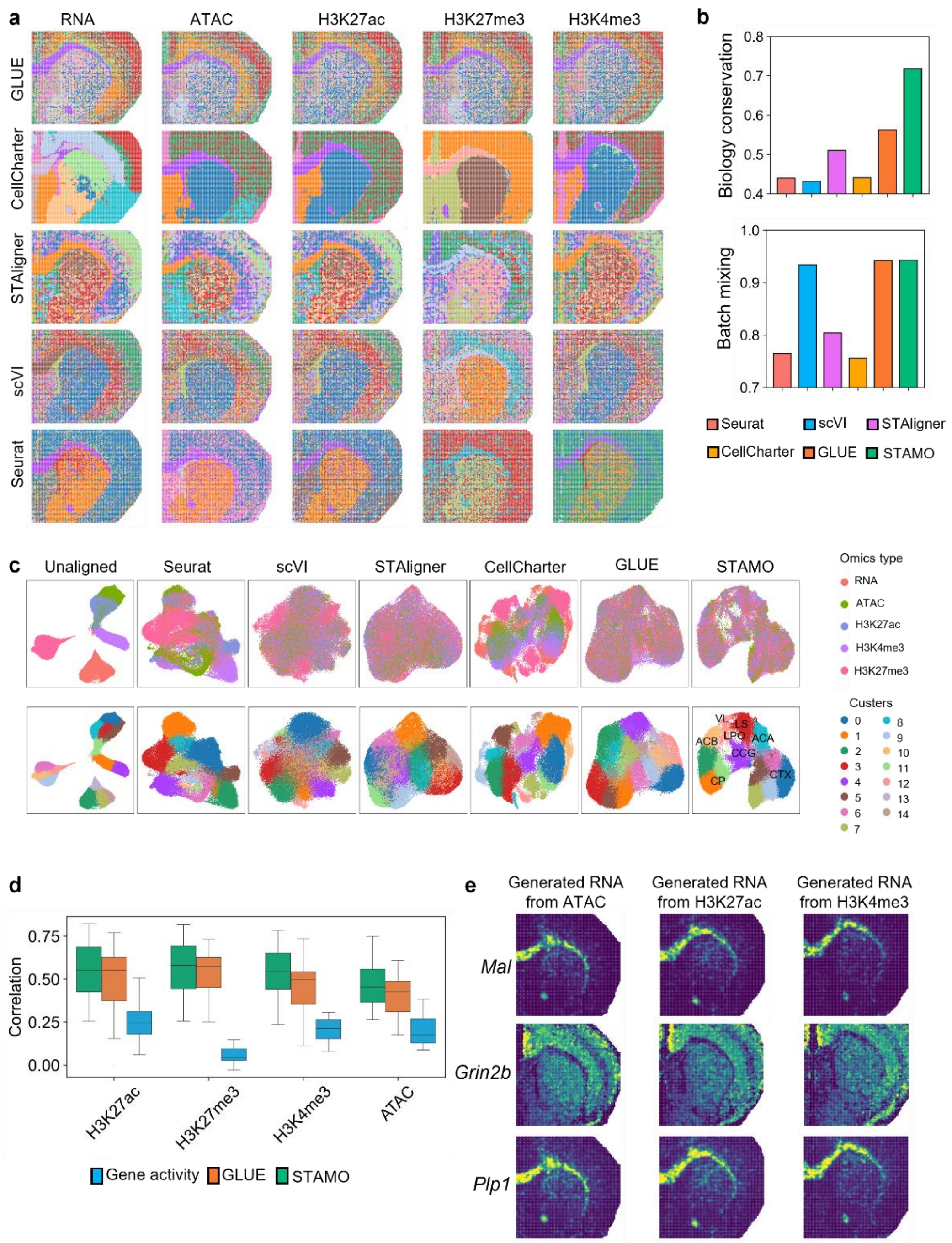


**Supplementary Fig. S9. a.** Spatial RNA, ATAC, H3K27ac, H3K27me3, and H3K4me3 of the P22 mouse brain, with spatial domains characterized by competing integration methods. **b.** Comparison of STAMO with competing methods using biology conservation and batch mixing score. **c.** UMAP plots of the embeddings from different integration methods. Spots are colored by omics type (top) and colored by cluster (bottom). **d.** Gene-wise correlation between the generated and measured gene expression, comparing STAMO with competing methods. **e.** Spatial heatmap of generated gene expression from ATAC, H3k27ac, and H3K4me3 data for CCG-specific marker genes *Mal*, *Grin2b*, and *Plp1*.

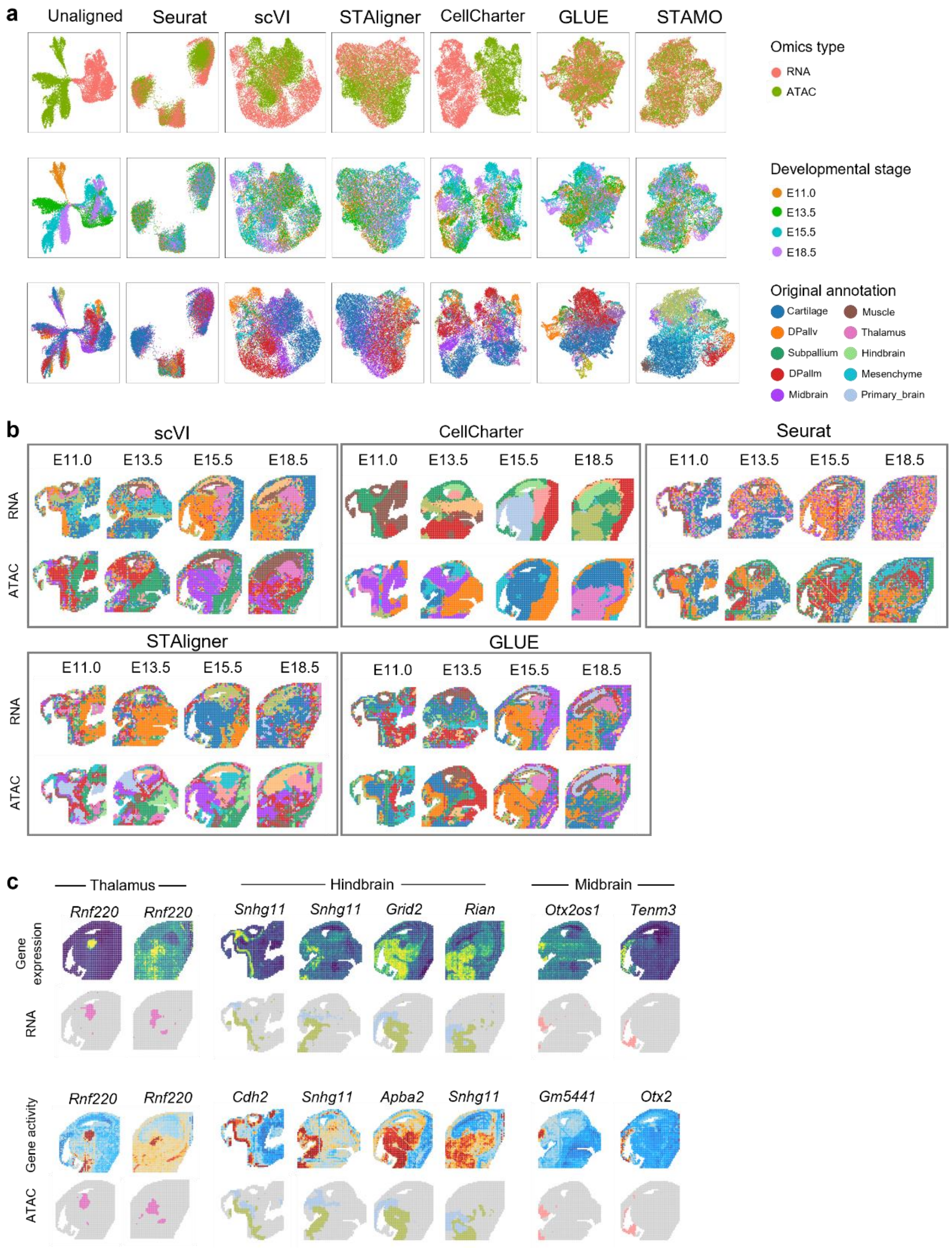


**Supplementary Fig. S10. a.** UMAP plots of embeddings of different integration methods colored by omics type (top), developmental stage (middle), and original tissue annotation (bottom). **b.** Spatial domains characterized by scVI, CellCharter, Seurat, STAligner, and GLUE. **c.** Spatial tissue substructures characterized by STAMO (bottom), and their corresponding marker genes (top).

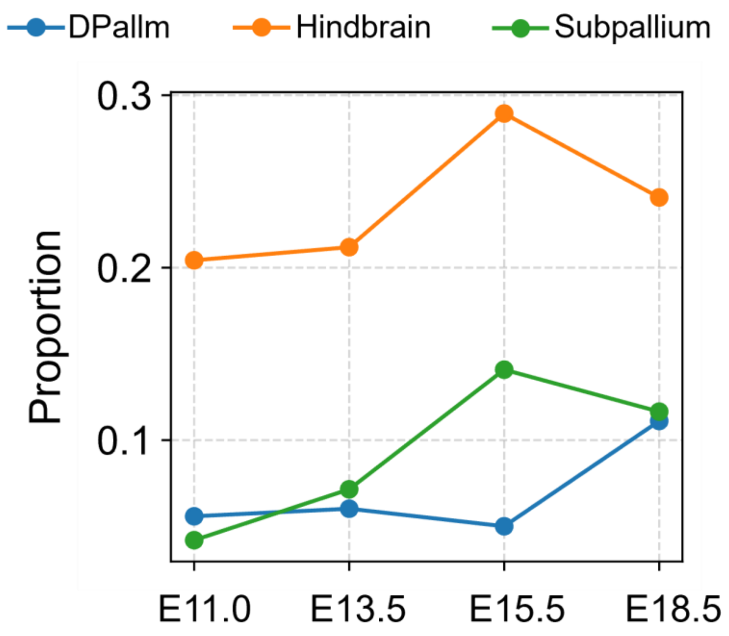


**Supplementary Fig. S11.** The proportions of DPallm, Hindbrain, and Suballium structures showing the developmental dynamics of tissue structures.

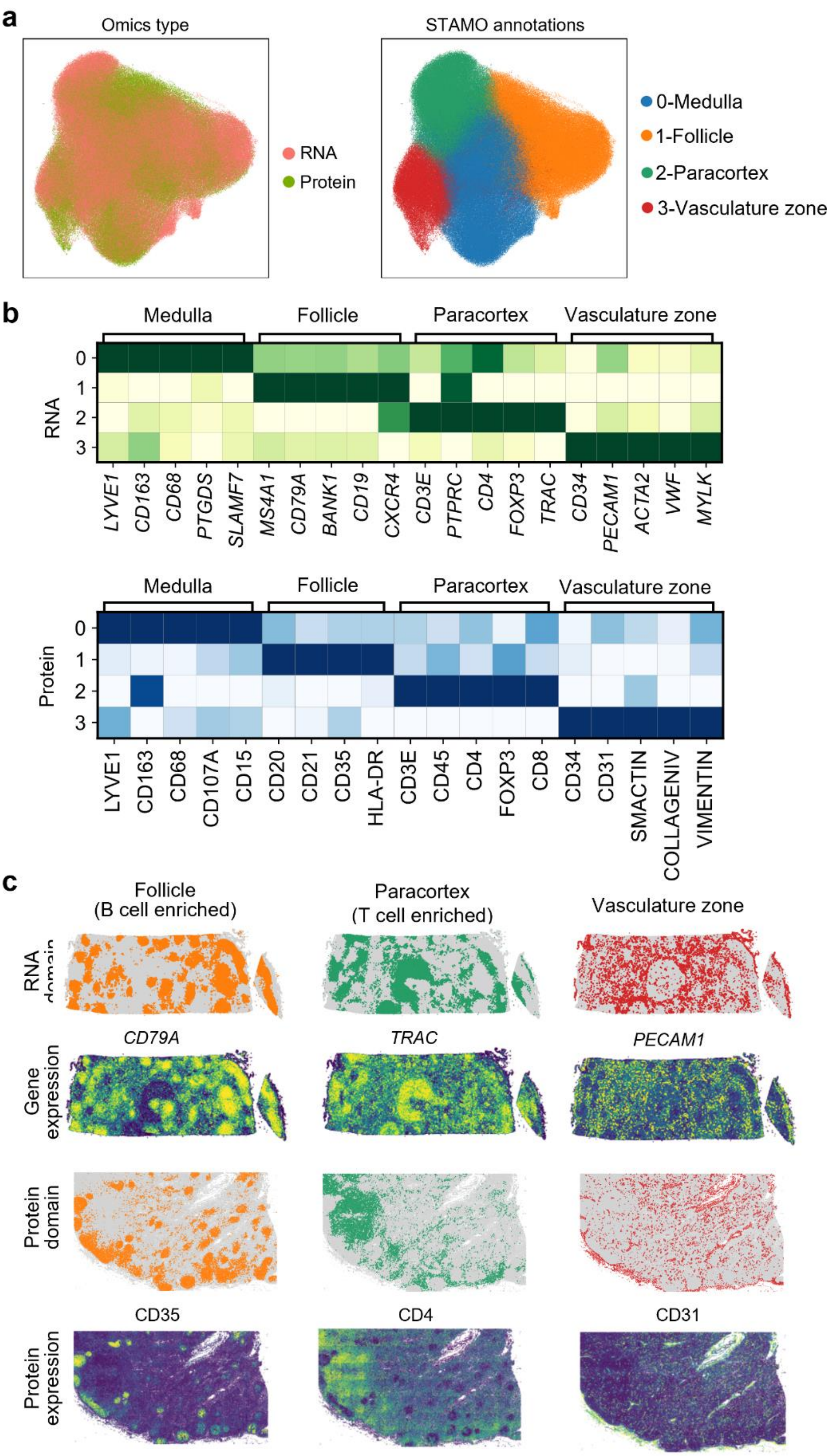

a
Omics type
RNA
Protein
STAMO annotations
0-Medulla
1-Follicle
2-Paracortex
3-Vasculature zone
b
Medulla
Follicle
Paracortex
Vasculature zone
RNA
0
1
2
3
LYVE1
CD163
CD68
PTGDS
SLAMF7
MS4A1
CD79A
BANK1
CD19
CXCR4
CD3E
PTPRC
CD4
FOXP3
TRAC
CD34
PECAM1
ACTA2
VWF
MYLK
Protein
LYVE1
CD163
CD68
CD107A
CD15
CD20
CD21
CD35
HLA-DR
CD3E
CD45
CD4
FOXP3
CD8
CD34
CD31
SMACTIN
COLLAGENIV
VIMENTIN
c
Follicle
(B cell enriched)
Paracortex
(T cell enriched)
Vasculature zone
RNA domain
Gene expression
CD79A
TRAC
PECAM1
Protein domain
Protein expression
CD35
CD4
CD31

**Supplementary Fig. S12. a.** UMAP plots of STAMO embeddings colored by omics (left) and tissue structure annotations (right). **b.** RNA and protein expression of region-specific markers (LYVE1 [1], CD163 [2], CD68 [3], PTGDS [4], SLAMF7 [5], MS4A1 [6], CD79A [7], BANK1 [8], CD19 [9], CXCR4 [10], CD3E [11], PTPRC [12], CD4 [13], FOXP3 [14], TRAC [15], CD34 [16], PECAM1 [17], ACTA2 [18], VWF [19], MYLK [20]) in STAMO's spatial domains. **c.** Spatial visualization of annotated tissue structures showing marker gene expression (top) and marker protein expression (bottom).

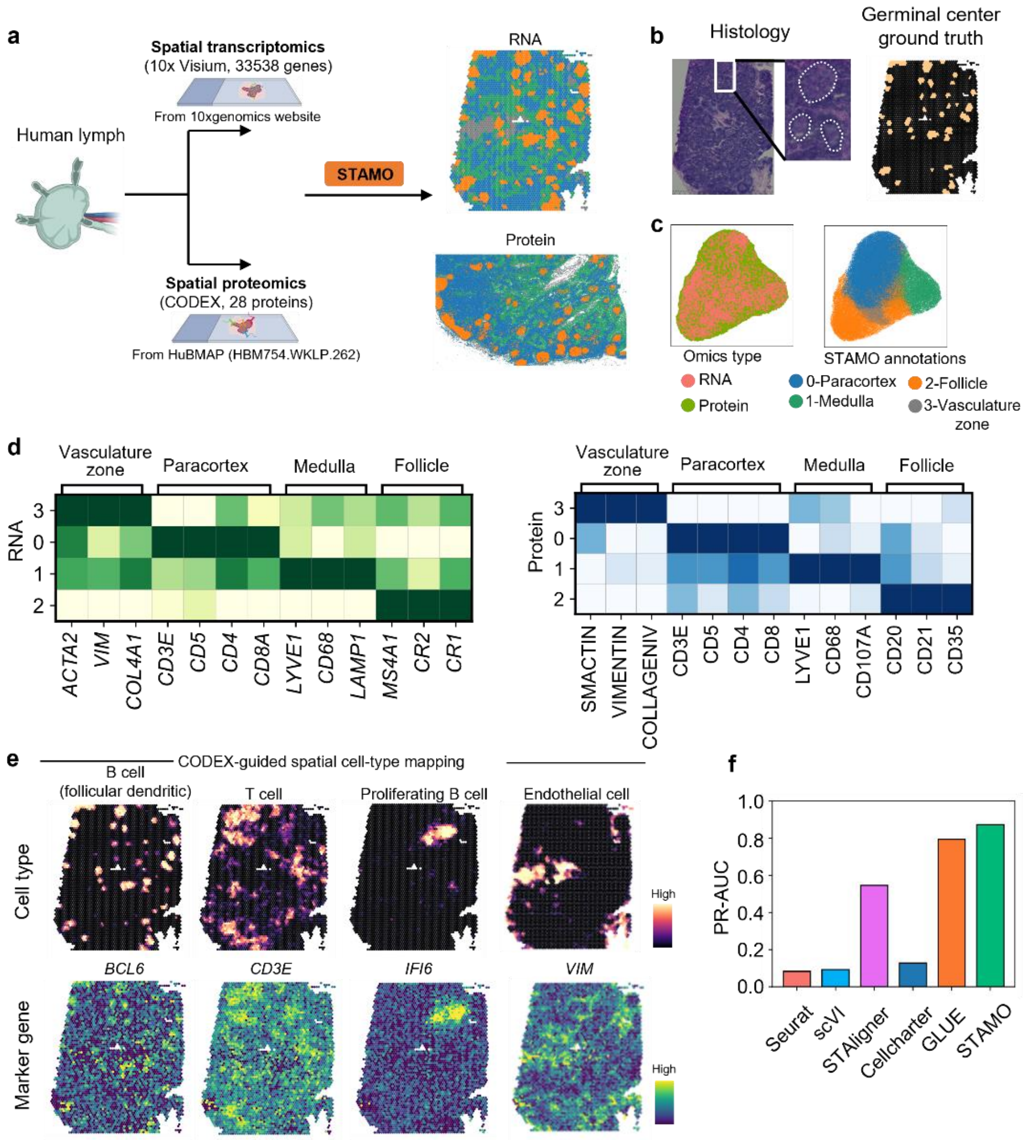


**Supplementary Fig. S13. Integration of human lymph spatial transcriptomics (Visium) and proteomics (CODEX) slices with limited overlap features. a.** Human lymph spatial transcriptomics slice produced by 10x Visium and spatial proteomics slice produced by CODEX (left). Visualization of identified spatial domains across slices characterized by STAMO (right). Created in https://BioRender.com. **b.** Annotation of germinal center (GC) regions based on paired histological images, with GC-positive locations highlighted in yellow and GC-negative locations shown in black. **c.** UMAP plots of STAMO embeddings colored by tissue structure annotations (left) and omics (right). **d.** Gene (left) and corresponding protein (right) expression of region-specific markers in STAMO's spatial domains. **e.** Spatial distributions of selected cell types transferred from CODEX slice (top) and their corresponding marker genes (bottom). **f.** Quantitative evaluation of mapping accuracy for follicular dendritic B cells, using the GC-positive and GC-negative locations in panel (**b.**) as reference. Precision-recall scores for different integration methods are shown.

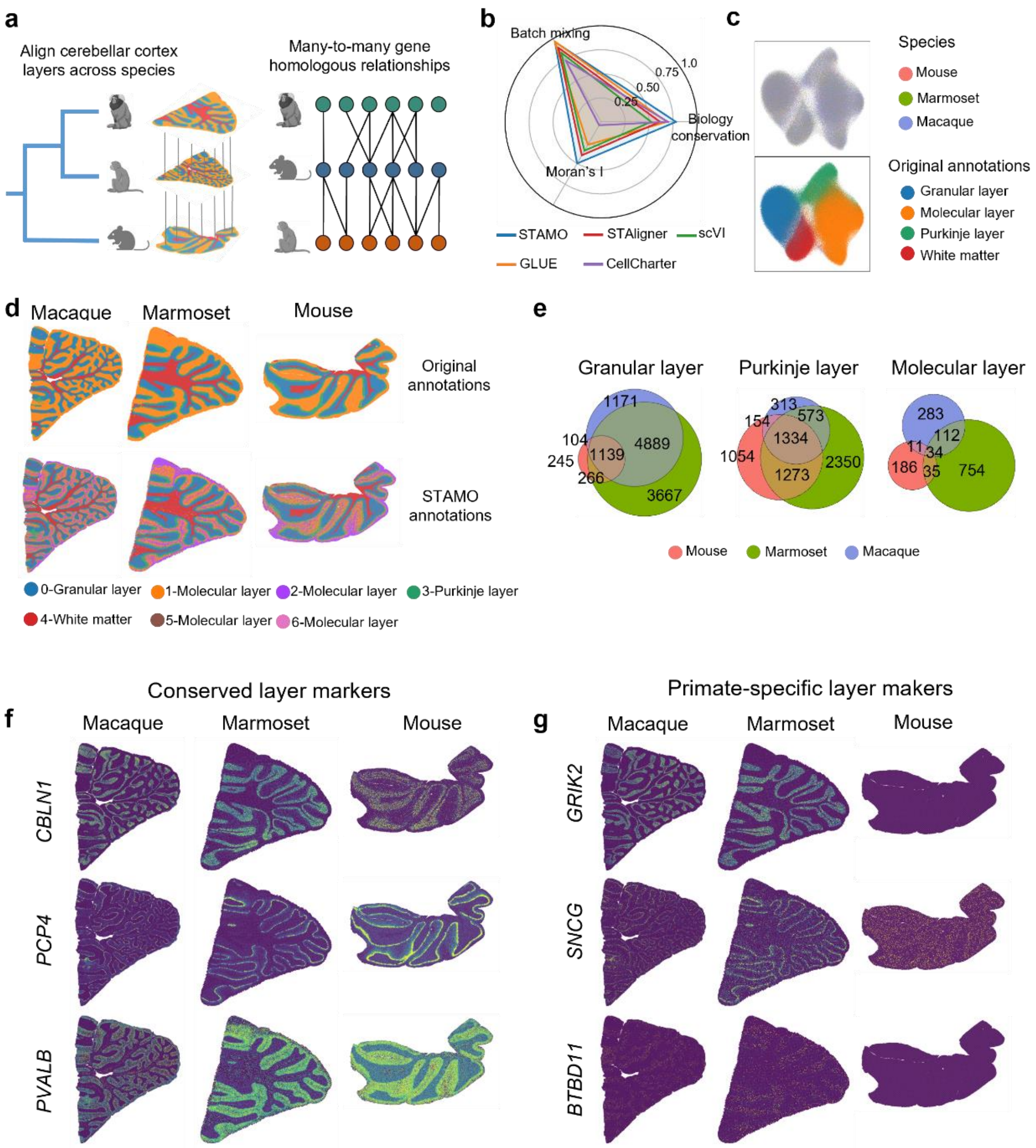


**Supplementary Fig. S14. Integration of cross-species spatial transcriptomics cerebellar cortex slices with many-to-many gene homologous relationships. a.** Stereo-seq slices from mouse, marmoset, and macaque with original cerebellar layer annotations (left) and many-to-many gene homology relationships (right). Created in https://BioRender.com. **b.** Moran's *I*, biology conservation, and batch mixing score of different integration methods. **c.** UMAP plots of STAMO embeddings colored by species (left) and raw annotations (right). **d.** Visualization of aligned spatial domains across slices characterized by STAMO. **e.** The number of conserved and divergent layer-enriched genes across species in the granular, Purkinje, and molecular layers. **f.** Spatial expression patterns of three known marker genes, *CBLN1*, *PCP4*, and *PVALB*, for the granular, Purkinje, and molecular layer, respectively. **g.** Spatial expression patterns of primate-specific layer-enriched genes for the granular, Purkinje, and molecular layer.

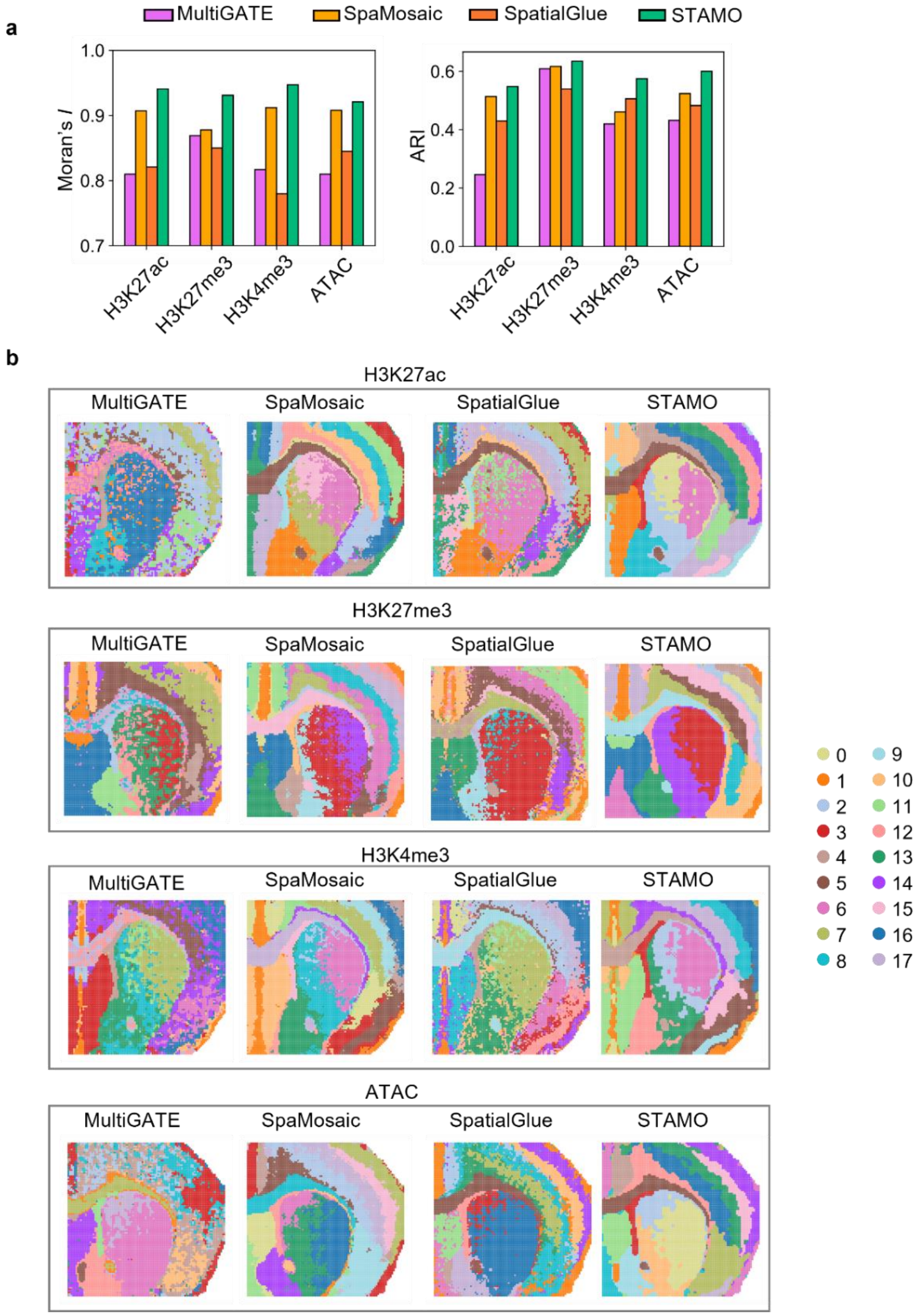


**Supplementary Fig. S15.** Comparison of STAMO with paired spatial integration method MultiGATE, SpaMosaic, and SpatialGlue. **a.** Quantitative evaluation of spatial coherence and clustering accuracy using Moran's *I* (left) and ARI (right) for each integration method. **b.** Spatial domain identification results of competing methods and STAMO on four paired P22 spatial multi-omics slices.

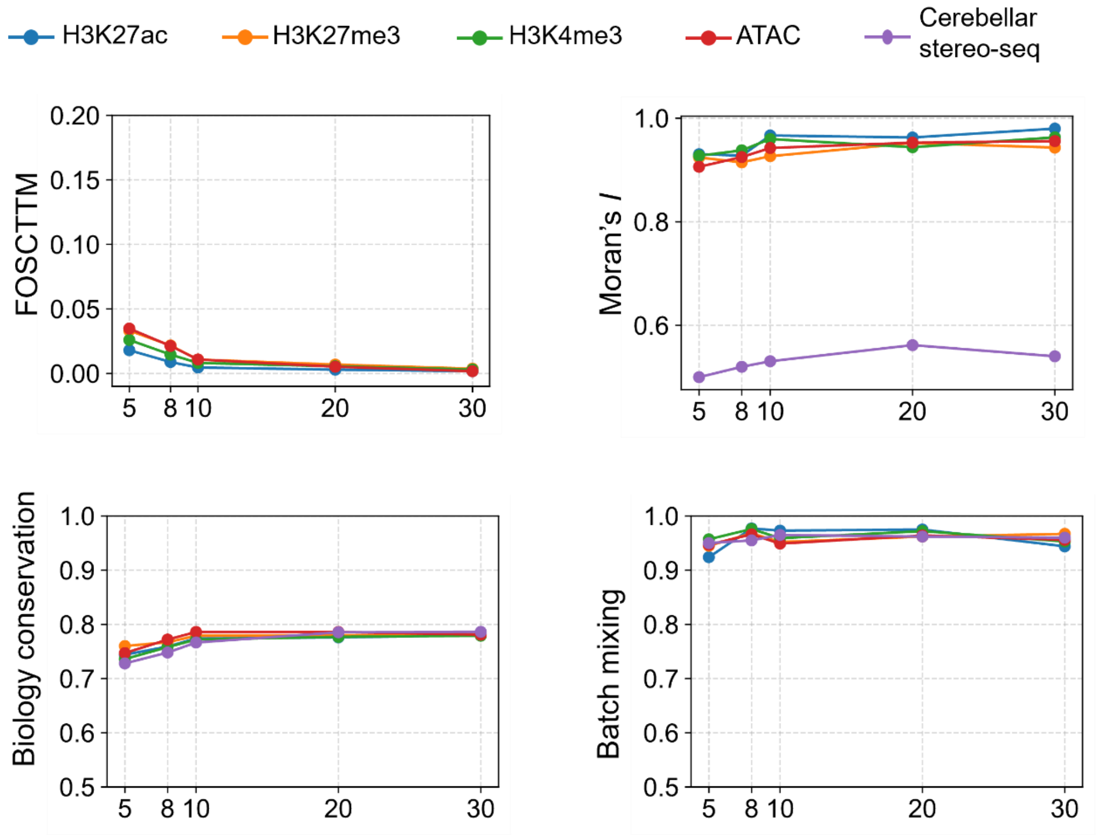


**Supplementary Fig. S16.** Integration performance of STAMO across different numbers of neighbors during spatial graph construction.

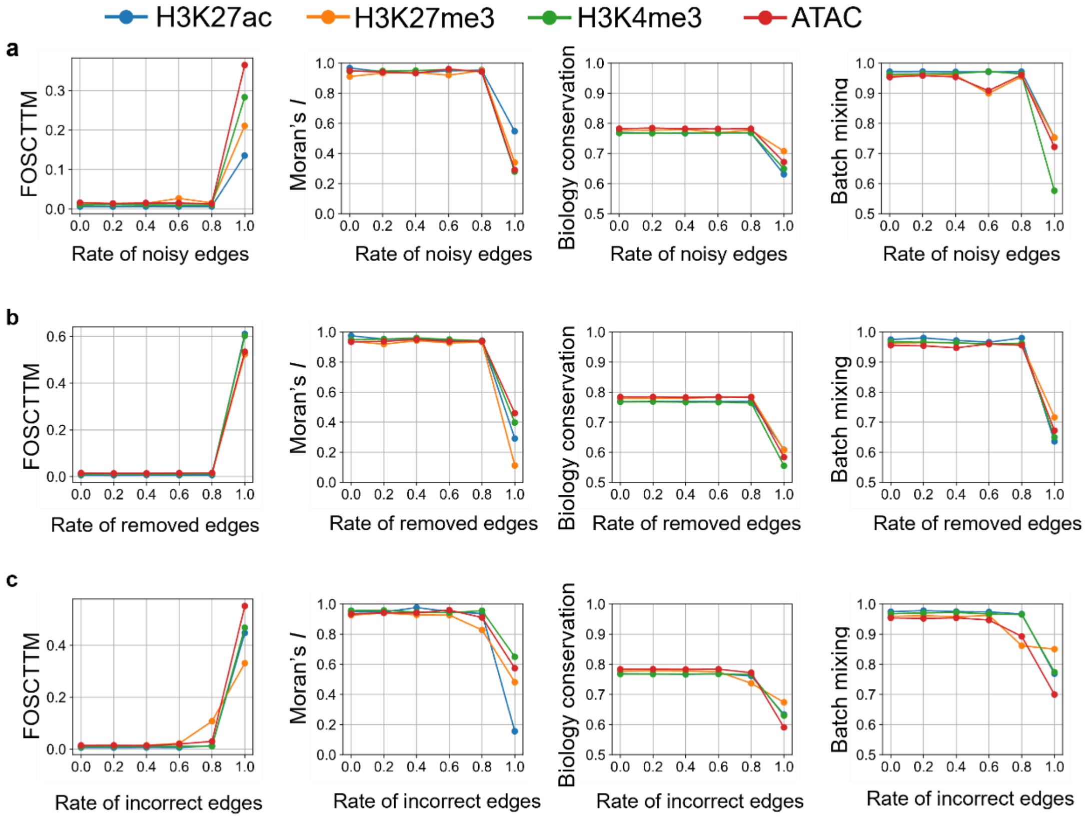


**Supplementary Fig. S17. Evaluation of STAMO's robustness on the prior feature graph. a.** Performance at increasing rates of noisy edges added to the prior graph. **b.** Performance at increasing rates of removal of correct edges from the prior graph. **c.** Performance at increasing rates of incorrect edges in the prior graph by simultaneously removing a proportion of true edges and replacing them with an equal number of nonexistent edges.

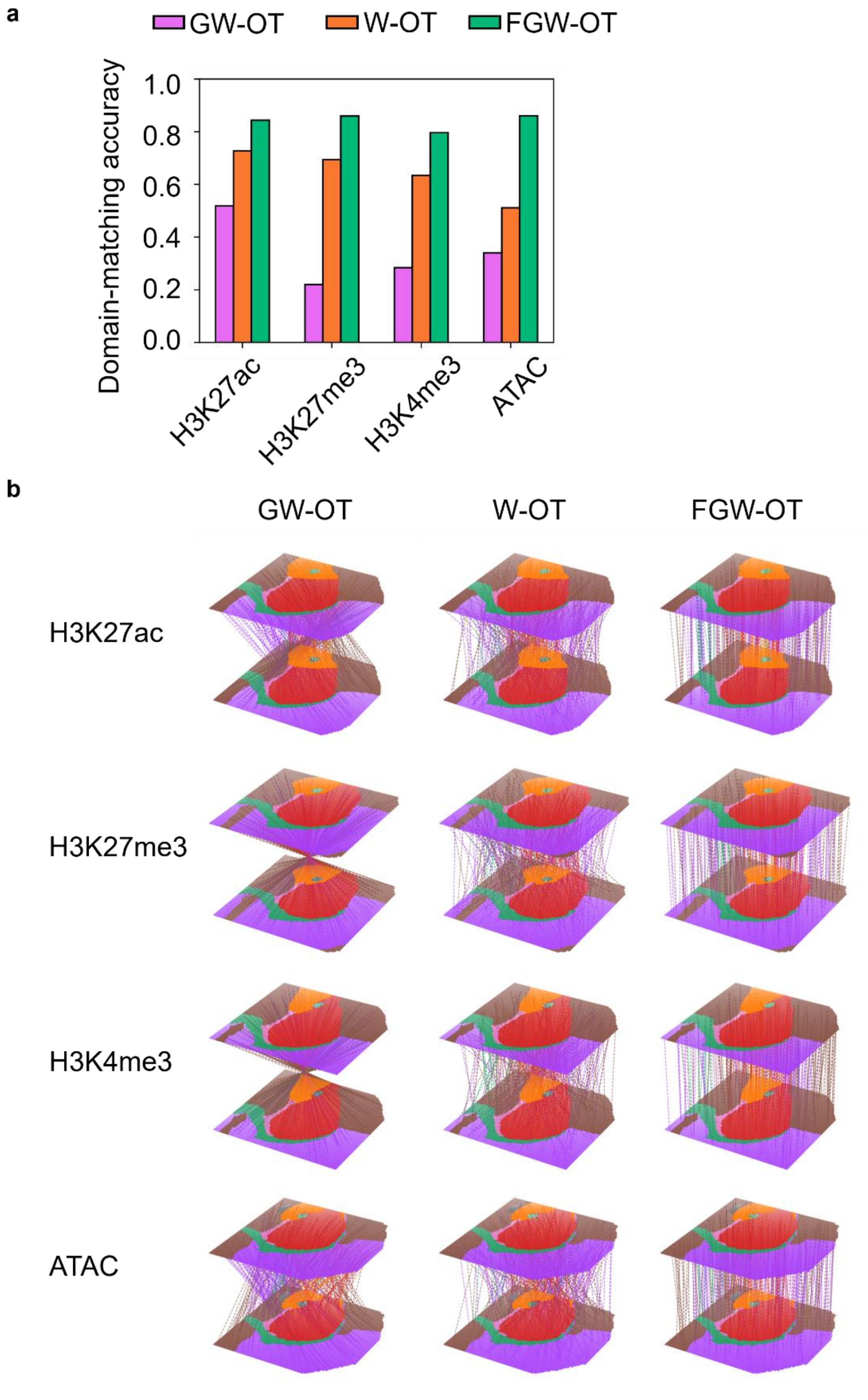


**Supplementary Fig. S18.** The anchor identification accuracy of different OT methods evaluated by spatial domain matching accuracy (**a**) and visualization of the alignment colored by manual annotations (**b**).

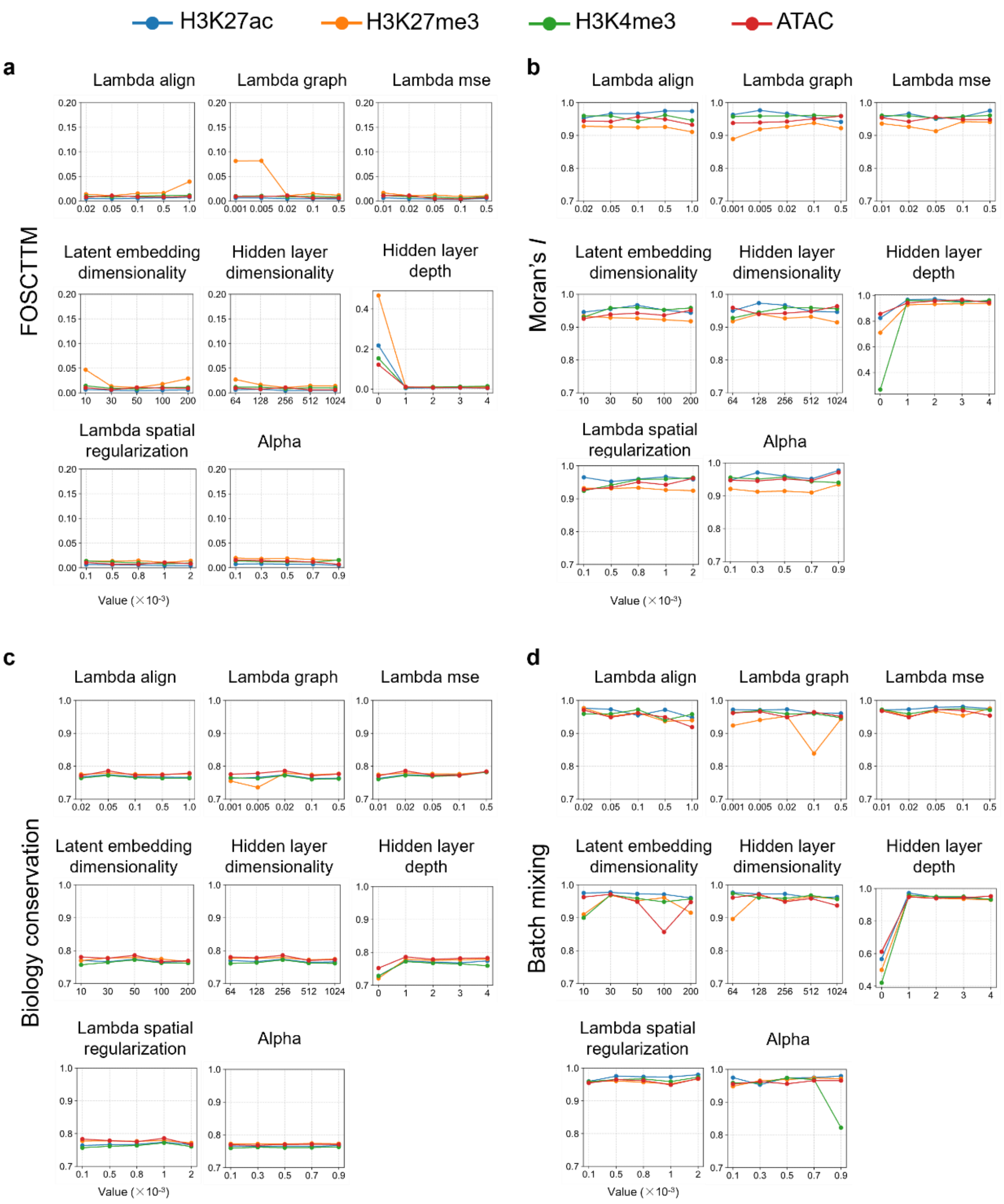


**Supplementary Fig. S19. Integration performance of STAMO under different hyperparameter settings. a.** Integration performance is quantified by FOSCTTM (**a.**), Moran's *I* (**b.**), biology conservation (**c.**), and batch mixing (**d.**). 'Lambda align' is the weight of the adversarial alignment ($\lambda_D$), 'Lambda graph' is the weight of the feature graph reconstruction loss $\lambda_{Gf}$, 'Lambda mse' is the weight of the FGW-OT alignment loss ( $\lambda_{mse}$ ), 'Lambda spatial regularization' is the weight of the spatial regularization loss ($\lambda_{Gs}$), 'Alpha' is the weight of trade-off parameter $\alpha$ of W-OT and GW-OT in FGW-OT.

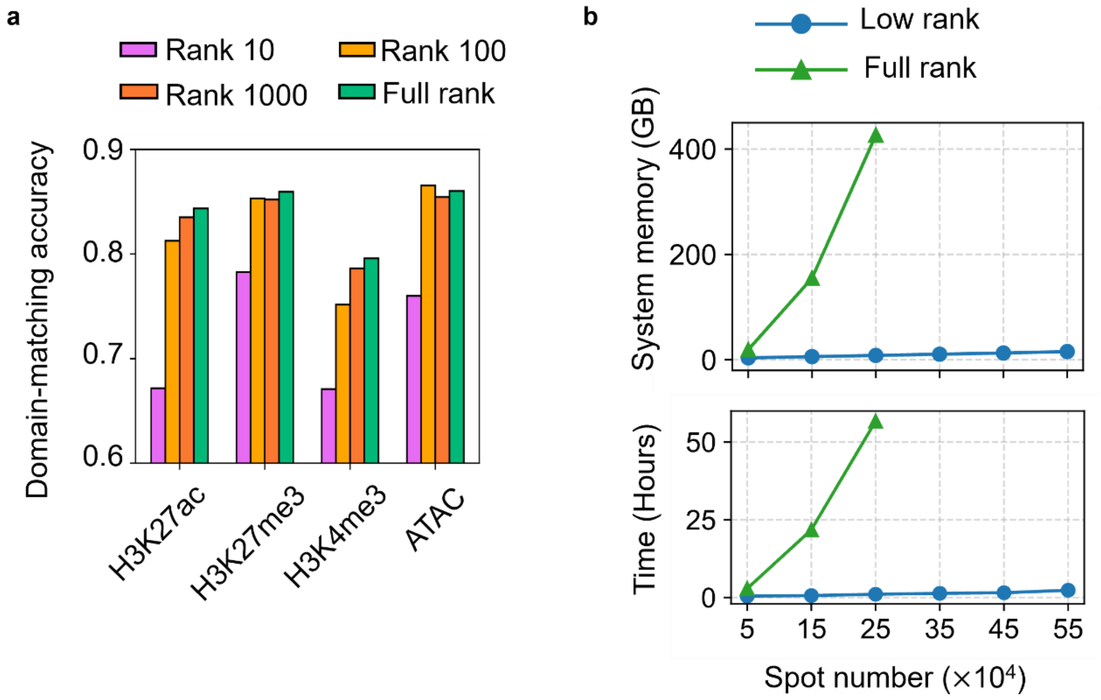


**Supplementary Fig. S20. a.** The spatial domain matching accuracy of FGW-OT under the full-rank and low-rank modes on P22 spatial epigenome-transcriptome mouse brain slices. **b.** Benchmark of peak memory consumption (top) and computation time (bottom) for increasing numbers of cells, subsampled from the human lymph node dataset.

#### Supplementary Tables

##### Supplementary Table 1. Technological advances of STAMO over existing spatial multi-omics integration methods

| | STAMO | SpatialGlue[21] | MISO[22] | CellCharter[23] | STAligner[24] |
|---|---|---|---|---|---|
| Spatial domain identification on unpaired data | ✓ | ✗ | ✗ | ✓ | ✓ |
| Cross-omics data generation on unpaired data | ✓ | ✗ | ✗ | ✗ | ✗ |
| Infer gene regulation network on unpaired data | ✓ | ✗ | ✗ | ✗ | ✗ |
| Infer positive and negative regulation simultaneously | ✓ | ✗ | ✗ | ✗ | ✗ |
| Cross-species alignment with non-one-to-one homologous gene mapping | ✓ | ✗ | ✗ | ✗ | ✗ |

**Supplementary Table 2.** Summary of the spatial multi-omics data used in this study.

| Tissue | Platform | Slice id | # of spots or cells | Related figures |
|---|---|---|---|---|
| **Mouse postnatal day 22 (P22) brain** | Spatial ATAC-RNA | ATAC-RNA | 9215 | Fig. 2-3<br>Fig. S1-5 |
| | Spatial CUT&Tag-RNA-seq | H3K27ac-RNA | 9370 | |
| | | H3K27me3-RNA | 9752 | |
| | | H3K4me3-RNA | 9548 | |
| **Mouse embryo** | MISAR-seq Spatial ATAC-RNA-seq | ATAC-RNA (E11.0) | 1263 | Fig. 4<br>Fig. S6 |
| | | ATAC-RNA (E13.5) | 1777 | |
| | | ATAC-RNA (E15.5) | 1949 | |
| | | ATAC-RNA (E18.5) | 2129 | |
| **Adult human lymph node** | CODEX (Protein) | N.A. | 188450 | Fig. 5<br>Fig. S7-8 |
| | 10x Xenium (RNA) | | 377897 | |
| | 10x Visium (RNA) | | 4039 | |
| **Mouse liver metastases** | Slide-DNA-seq | N.A. | 24578 | Fig. 6 |
| | Slide-RNA-seq | N.A. | 25318 | |
| **Macaques, marmosets, and mice cerebellar cortex** | Stereo-seq | Macaque1_T88 | 345943 | Fig. S9 |
| | | Marmoset1_T502 | 97791 | |
| | | Mouse2_T353 | 92163 | |
